# Loss of Cerebral Autoregulation After Stroke Drives Abnormal Perfusion Patterns

**DOI:** 10.1101/2025.10.03.680208

**Authors:** Chryso Lambride, Mohamad El Amki, Susanne Wegener, Franca Schmid

## Abstract

Cerebral autoregulation stabilizes cerebral blood flow (CBF) in response to variations in blood pressure, primarily through dynamic adjustments in arterial caliber. However, the impact of autoregulation mechanisms on local vascular responses and microcirculatory perfusion remains poorly understood. Given that impaired autoregulation is common in pathologies such as ischemic stroke, quantifying vessel-level responses can help to inform clinical strategies. Here, we present a novel *in silico* model that incorporates static myogenic and endothelial regulatory mechanisms to simulate CBF in large microvascular networks derived from realistic surface vasculatures of mice with and without leptomeningeal collaterals (LMCs). We assessed the role of autoregulation mechanisms under three conditions: i) healthy autoregulation, ii) ischaemic stroke and reperfusion with altered vascular reactivity, and iii) chronic autoregulation dysfunction after stroke. For healthy autoregulation, our model reproduced the classic static autoregulation curve and identified the dominant contribution of surface arteries in buffering pressure changes. Networks with lower descending arteriole density exhibited more extensive arterial dilations, reflecting the topological influence on vessel diameter changes. During reperfusion after stroke, we investigated the interplay between LMCs and parameters governing the vasoreactive response. While LMCs play a crucial role in maintaining residual perfusion in adjacent regions during MCA occlusion, our results suggest that their extent alone does not substantially influence perfusion after recanalization. Instead, alterations in myogenic reactivity emerged as the key contributor to hyperperfusion, underscoring the importance of regulatory mechanisms in determining reperfusion outcomes. To mimic chronic autoregulatory dysfunction, we progressively impaired regulatory capacity in arteries on the middle cerebral artery (MCA) side, reflecting the territory previously affected by stroke. The loss of autoregulation in proximal vessels significantly disrupts capillary perfusion and alters the autoregulation curve. To our knowledge, this is the first *in silico* study to explore cerebral autoregulation at the microvascular level under both physiological and pathological conditions. Our findings provide mechanistic insight into how individual vessel behavior shapes global flow regulation and highlight myogenic tone as a potential therapeutic target to reduce reperfusion-related complications in stroke.

## Introduction

Cerebral autoregulation is a key physiological process that maintains relatively constant cerebral blood flow (CBF) across a range of blood pressure variations, primarily through adjustments in arterial diameter^1,2^. This regulation is essential for healthy brain function and relies on a complex interplay of vascular control mechanisms. Although the underlying mechanisms are not yet fully elucidated, current evidence points to modulation of vascular smooth muscle (VSM) tone as the primary driver of CBF regulation^3^. Among the cerebral vessels, arteries stand out as key regulators, providing about 50% of total vascular resistance and containing a high density of VSM cells (VSMCs) capable of adjusting tone in response to pressure changes^4,5^.

Both VSMCs and endothelial cells contribute critically to this regulation^2,6–9^, largely due to their mechanosensitive properties, which allow them to respond to changes in transmural pressure and shear stress^7,9^. The endothelial (flow-mediated) response is triggered by increased shear stress, leading to VSM relaxation and subsequent vasodilation^2,10^, whereas the myogenic response involves intrinsic VSM constriction in response to elevated intravascular pressure^2,3^. This myogenic behavior is considered independent of neural, metabolic, or hormonal influences^3^. The net effect on vascular tone reflects the balance between these opposing mechanisms and is mediated *via* the degree of VSM phosphorylation^11,12^.

While many studies have characterized cerebral autoregulatory mechanisms in detail^4,5,13–17^, the contribution of individual arteries and the resulting flow changes at single-vessel resolution remain largely unexplored. Quantifying these responses *in vivo* is particularly challenging due to the concurrent activity of multiple physiological processes, which makes it difficult to isolate the effects of specific mechanisms. To address this, we developed a static autoregulation model incorporating pressure- and flow-dependent responses representing myogenic and endothelial regulation. This *in silico* framework is applied to large, semi-realistic microvascular networks reconstructed from mouse brains, enabling single-vessel resolution under both physiological and pathological conditions.

During healthy autoregulation, important questions remain about how individual arterial segments contribute to CBF regulation at the microvascular level. Specifically, it is unclear whether autoregulation involves a uniform vascular adjustment or reflects a hierarchical pattern of responsiveness across the arterial vasculature. If such a hierarchy exists, the next question is how it might be characterized, potentially through correlations with topological features such as vessel size or spatial location within the network. Moreover, studies that quantify how structural differences across brain regions or individuals, such as penetrating arterial tree density, affect the network’s ability to maintain stable perfusion under pressure variations are still lacking.

Quantifying these spatial differences in cerebral autoregulation is particularly relevant in disease states where vascular tone regulation becomes disrupted. Dysfunction of autoregulation has been implicated in a range of pathological conditions, including ischaemic stroke^18–20^, traumatic brain injury^21,22^, orthostatic hypotension^23^, and neurodegenerative diseases such as Alzheimer’s disease^24,25^. In the context of ischaemic stroke, typically caused by a blood clot obstructing a major cerebral artery^26^, experimental studies have primarily focused on the impact of stroke on arteries within the affected region. A key finding suggests that during ischaemia followed by reperfusion, these arteries might lose their ability to constrict in response to elevated intravascular pressure, a hallmark of disrupted myogenic response^14,27^.

Clinical studies have examined changes in cerebral autoregulation across the acute (<48 hours), subacute (48 hours to 7 days), and chronic (>7 days) phases of ischaemic stroke^19^. Cerebral autoregulation can be evaluated as either a static or dynamic phenomenon. Notably, in the acute ischaemic stroke setting where blood pressure manipulation carries significant risk, static assessments are used cautiously to avoid aggravating ischaemic injury. Hence, limited clinical data on static autoregulation is available^19,28–32^. Direct comparisons across studies are challenging due to substantial heterogeneity in study design, including differences in timing of assessments, stroke subtypes and severities, and the metrics and methodologies used to quantify autoregulation^19,29,31^. Such variability likely contributes to the mixed findings reported in the literature, especially during the acute phase. While findings in the acute phase remain inconsistent, more robust evidence points to impaired autoregulation in the subacute phase, with partial or full recovery occurring during the chronic phase^19^. In some patients, particularly those with more severe strokes or poorer functional outcomes, these impairments may persist beyond the subacute phase^19,29^. When autoregulation is impaired, arteries no longer actively respond to changes in pressure but instead behave passively, leading to potentially harmful variations in cerebral perfusion even with small blood pressure fluctuations. Despite its clinical importance, how chronic autoregulation dysfunction affects perfusion across different vessel types remains largely unexamined.

Interestingly, during the acute phase, alterations in autoregulatory mechanisms might cause different reperfusion dynamics, thus determining the post-stroke outcome^27,30,33–36^. Recent *in vivo* studies have shown that reperfusion profiles following middle cerebral artery (MCA) occlusion tend to be correlated with the extent of leptomeningeal collaterals (LMCs)^37^. LMCs, which are anastomotic connections between major cerebral arteries, provide alternative pathways for blood flow to sustain perfusion to under-perfused regions during arterial occlusion. Their extent varies widely among individuals, and richer collateral networks are generally associated with reduced infarct volumes and better clinical outcomes^26,38,39^. Specifically, mice with poor collateralization exhibit rapid and excessive hyperaemic reperfusion in proximal MCA territories, whereas those with abundant LMCs undergo a more gradual and controlled restoration of flow. These contrasting reperfusion profiles have led to the hypothesis that poor collateral networks result in ischaemia-induced endothelial damage, thereby impairing vasoregulatory capacity in the recanalized MCA segment. In contrast, rich collateral networks may sustain minimal perfusion during occlusion, preserving endothelial function and enabling more physiologically regulated reperfusion upon recanalization^37^. However, this hypothesis has not yet been systematically tested. Particularly, it remains unclear how the extent of LMCs influences perfusion in the proximal MCA territory during ischaemic stroke and whether collateralization serves as the principal determinant of these divergent reperfusion responses.

Computational models offer a powerful alternative, enabling the controlled investigation of individual factors and their contributions to cerebral haemodynamics. Over the past decades, numerous computational models have been developed to describe both static and dynamic cerebral autoregulation^12,40–48^. Most of these employ a compartmental approach, in which vascular territories or vessel segments are simplified and grouped into lumped compartments. While this framework offers computational efficiency and yields insights into global autoregulatory behavior, it drastically simplifies and overlooks the structural and functional heterogeneity of the microvascular network. In particular, compartmental models do not explicitly represent individual vessels or their morphological attributes, limiting their ability to investigate mechanisms at a single-vessel resolution. As such, they cannot account for the spatial heterogeneity and local feedback mechanisms that are critical for understanding blood flow regulation at the microvascular level during both normal and disease conditions.

Recent advances have attempted to overcome these limitations. Notably, work by Daher *et al*.^49^ introduced a network-based modelling approach to dynamic cerebral autoregulation, incorporating both myogenic and endothelial responses. Additionally, Esfandi *et al*.^50^ developed a static autoregulation model to assess depth-dependent vascular contributions to pressure regulation and functional hyperemia using a simplified microvascular network. Their findings underscored the dominant role of small-diameter arterioles, particularly penetrating and precapillary vessels, in controlling capillary perfusion. However, their model did not explicitly incorporate distinct myogenic and endothelial control mechanisms to quantify microvascular responses under varying pathological conditions.

Here, we present a novel static autoregulation model applicable at the level of individual arteries within large, semi-realistic microvascular networks, incorporating both myogenic and endothelial control mechanisms. Our model is adapted from the dynamic autoregulation framework proposed by Daher *et al*.^49^, which we extended and reformulated to capture static autoregulatory behaviour. Using our *in silico* framework, we address key research questions related to healthy and impaired cerebral autoregulation following ischaemic stroke. Under healthy autoregulation conditions, we quantify vascular responses to changes in mean arterial pressure (MAP) and evaluate how the density of penetrating arterial trees influences autoregulatory capacity. We also simulate scenarios that mimic chronic autoregulatory dysfunction after stroke to quantify how the severity of impairment modifies the classical static autoregulation curve and affects capillary-level perfusion. Finally, we apply our model during stroke-reperfusion to investigate the role of LMCs in shaping reperfusion dynamics and to examine the changes in control mechanisms required for reproducing the hyperperfusion patterns observed in *in vivo* experiments.

## Methods

### Microvascular Networks

Our simulations have been performed in four large semi-realistic microvascular networks (MVNs) previously generated by Epp *et al*.^51^. These MVNs were derived from two mouse strains that differ in collateral extent and the density of arterial penetrating trees. These variabilities make our MVNs particularly well-suited for investigating how autoregulation is achieved in different microvascular architectures. A detailed description of the generation and structure of these MVNs is provided in^51^; here, we briefly summarise the key steps, as shown in Figure 1A.

**Figure 1.**
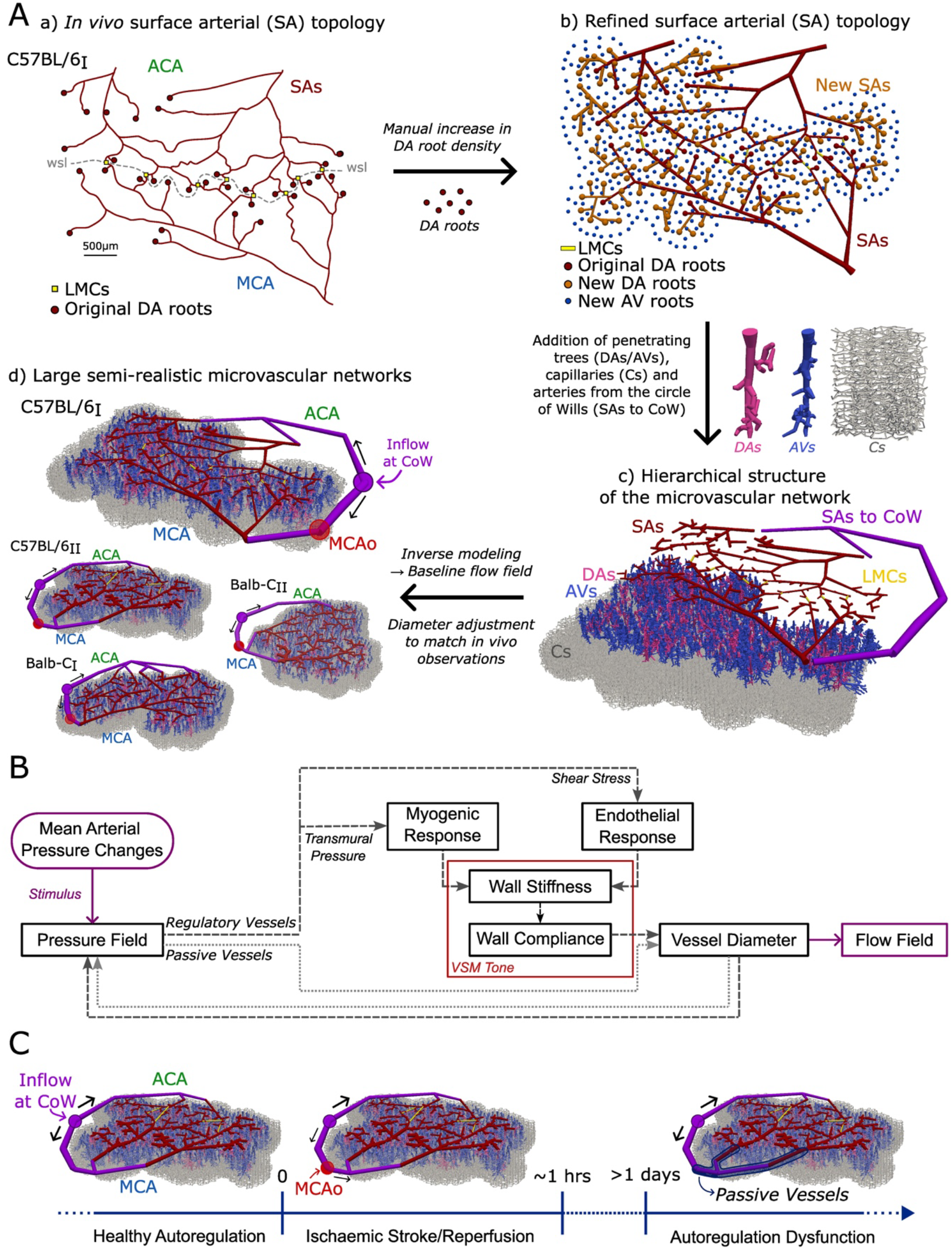
Generation of large semi-realistic microvascular networks and overview of the autoregulation model. **(A)** Stepwise generation of large semi-realistic microvascular networks. (a) *In vivo* surface arterial (SA) topology of the C57BL/6I mouse showing middle cerebral artery (MCA) and anterior cerebral artery (ACA) branches connected by leptomeningeal collaterals (LMCs, yellow squares) across the watershed line (wsl); original DA root points are shown in red. (b) Refined SA topology with increased DA root density (orange circles), distributed ascending vein (AV) roots (blue circles) and newly added SAs (orange lines). (c) Hierarchical network structure including DAs, AVs, and an artificial capillary bed (Cs). Additional SAs are added to connect MCA and ACA territories to a common inflow node representing the Circle of Willis (CoW). (d) Large semi-realistic microvascular networks generated from C57BL/6 and Balb-C mice, showing inflow locations at CoW (purple circles) and MCA occlusion (MCAo) sites (red circles). **(B)** Schematic representation of the autoregulation model. The flow chart for active and passive vessels are depicted by dashed and dotted lines, respectively. **(C)** Schematic timeline of simulated autoregulatory conditions including (i) healthy conditions, (ii) ischaemic stroke followed by reperfusion (~1 hour), and (iii) chronic autoregulation dysfunction (>1 day post-occlusion).

The MVNs were reconstructed from *in vivo* surface arterial (SA) networks of C57BL/6 (LMC-rich) and Balb-C (LMC-poor) mice acquired by two-photon imaging. Each network covers a 3.5 × 3.5 mm^2^ region of the cortical surface in the whisker and hindlimb areas, capturing vessels from both the anterior cerebral artery (ACA) and middle cerebral artery (MCA) territories (MCA-M4/M5). Descending arteriole (DA) density has been increased to match strain-specific values obtained from *in vivo* quantifications, with C57BL/6 mice exhibiting higher densities than Balb-C mice^52^. To complete the cortical microvascular architecture, ascending venules (AVs) were added and connected to DAs by an artificial capillary bed. Additional arteries (SAs-to-CoW) are included to connect the MCA- and ACA-sided SAs to a common inflow vertex, coming from the circle of Willis (CoW). In agreement with experiments^37^, occlusion for stroke simulations is induced at the MCA-M2 bifurcation located along the MCA-sided SAs to CoW arteries. To reduce uncertainty in vessel diameters and to guarantee agreement with *in vivo* flow characteristics, an inverse modeling approach was applied to all networks^51,53^.

### Blood flow model

The cerebral microvasculature forms a dense, highly interconnected network of blood vessels, which can be modelled as a graph composed of edges and vertices. Each edge represents a blood vessel, and each vertex denotes a junction connecting at least two vessels. As commonly employed for microvascular blood flow^54–60^, we computed flow characteristics using Poiseuille’s law. This approach is appropriate for the microcirculation, because both the Reynolds number (Re < 1) and the Womersley number (Wo < 1) are low, ensuring laminar and quasi-steady flow.

The blood flow rate *q*_*ij*_ through each edge *e*_*ij*_ was computed as:

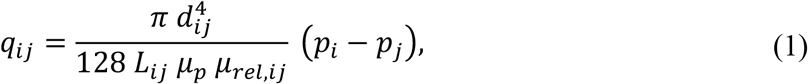

where *d*_*ij*_ and *L*_*ij*_ are diameter and length of the vessel associated with edge *e*_*ij*_. *μ*_*p*_ is plasma viscosity, and *μ*_*rel,ij*_ is the relative apparent viscosity accounting for the Fåhræus– Lindqvist effect. *p*_*i*_ and *p*_*j*_ are the pressure values at the source and target vertex *v*_*i*_ and *v*_*j*_ of edge *e*_*ij*_,We accounted for the impact of red blood cells on flow resistance by incorporating the empirical viscosity model developed by Pries and Secomb^61^ (*in vitro* version), which estimates *μ*_*rel,ij*_ based on vascular diameter and tube hematocrit *H*_*t,ij*_. Given our focus on the global flow field rather than capillary specific perfusion, we neglected phase-separation effects at bifurcations and assumed a uniform hematocrit *H*_*t,ij*_ = 0.3 throughout the network. We define a flow balance equation (continuity equation) *g*_*i*_ for each vertex *v*_*i*_ as:

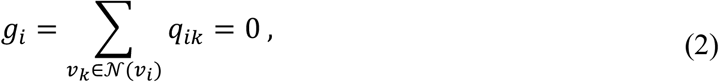

where *𝒩*(*v*_*i*_) is the set of all neighboring vertices connected to vertex *v*_*i*_. By combining Eqs. (1) and (2), a system of linear equations is derived. Solving this system with appropriate boundary conditions yields the pressure distribution across the entire network, from which flow rates are computed.

The MVNs contain a single inflow vertex, located at the most upstream SA connected to the CoW, and multiple outflow vertices corresponding to the roots of AVs. Pressure boundary conditions of 100 mmHg (inflow) and 10 mmHg (outflow) were applied^51^. The inflow pressure at the CoW was assumed to represent the MAP.

### Static cerebral autoregulation model

We modeled blood vessels as elastic tubes whose diameters adapt to changes in pressure either *via* passive and active mechanisms. When vessels respond passively, they dilate with rising pressure or collapse with falling pressure, leading to pressure-dependent changes in CBF. Unlike the passive response, active behavior arises from cerebral autoregulatory mechanisms, which adjust VSM tone to maintain a stable CBF despite variations in MAP. These active responses allow the blood (regulatory) vessels to constrict or dilate to counteract pressure changes and preserve constant perfusion. In our model, we distinguish between passive (non-regulatory) and regulatory vessels. Passive vessels were assumed AVs and Cs, while regulatory vessels were considered all arteries including SAs to CoW, SAs and DAs, unless stated otherwise.

To model the passive responses, we used a pressure–area relationship derived from linear elastic theory (neglecting viscoelastic effects), that describes how transmural pressure alters the vessel cross-sectional area^62^. While many linear and non-linear relationships have been proposed^62–65^, we employed a widely used non-linear formulation expressed as follows^64^:

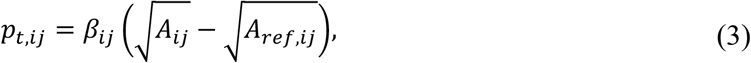

where *p*_*t,ij*_ = *p*_*ij*_ − *p*_*ext,ij*_ is the transmural pressure of each vessel, computed as the difference between the intraluminal *p*_*ij*_ and extravascular pressure *p*_*ext,ij*_. The intraluminal pressure is the average vessel pressure *p*_*ij*_ = 0.5(*p*_*i*_ + *p*_*j*_ / and *p*_*ext,ij*_ = 0 *mmHg* is assumed^41,49,62^. *A*_*ij*_ is the current cross-sectional area of the vessel and *A*_*ref,ij*_ indicates the reference vascular cross-sectional area at zero transmural pressure. *A*_*ref,ij*_ (*r*_*ref,ij*_, reference radius) of all vessels were calculated, using the diameters and pressure field at baseline conditions, based on the linear elastic theory:

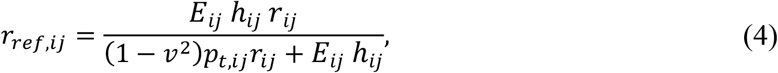

where *r*_*ij*_ is the vessel radius, *E*_*ij*_ indicates Young’s Modulus, *h*_*ij*_ is vascular wall thickness, and *v* is Poisson’s ratio, where *v* = 0.5 assuming that the vessel wall material is incompressible. The parameter *β*_*ij*_ is characterized by constant material properties and governs passive responses to pressure *via* Eq. (3). The coefficient of proportionality *β*_*ij*_ is a material parameter defined by^64^:

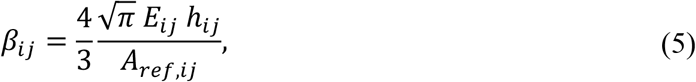

Therefore, when vessels behave passively, diameters *d*_*ij*_ were calculated by combining Eqs. (3) and (5):

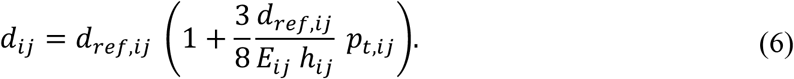

In contrast, regulatory vessels can actively adjust their wall vessel compliance through autoregulatory mechanisms, thereby modifying the pressure–area relationship^11,49^. To capture this behavior, we formulated a method to actively modulate Eq. (3) based on the core principles of cerebral autoregulation, by controlling the level of VSM tone through changes in compliance (or equivalently, stiffness). Compliance *C*_*ij*_ can be defined as the change in arterial volume *V*_*ij*_ relative to the change in pressure *p*_*t,ij*_ ^49^:

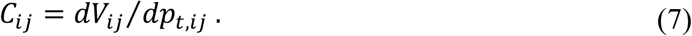

Applying the compliance definition (Eq. (7)) to the pressure-area relationship in Eq. (3) and assuming a constant area along the vessel such that *V*_*ij*_ = *L*_*ij*_ *A*_*ij*_, compliance *C*_*ij*_ can be also expressed in terms of *β*_*ij*_ as:

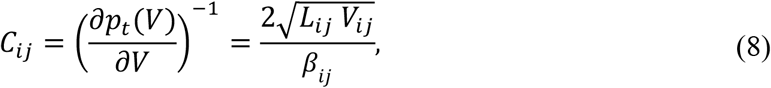

Accordingly, baseline compliance *C*_*o,ij*_ can be written as a function of the baseline radius *r*_*o,ij*_ and reference radius *r*_*ref,ij*_ as:

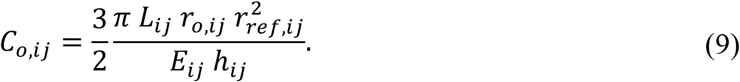

To simulate active regulation, we incorporated a quasi-steady feedback model that captures both myogenic and endothelial responses. Following Daher et al.^49^, the relative vessel wall stiffness *k*_*ij*_ is given by:

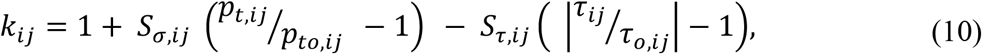

where *S*_*σ,ij*_ and *S*_*τ,ij*_ are sensitivity factors for myogenic and endothelial mechanisms, respectively. *S*_*σ,ij*_ was set higher than *S*_*τ,ij*_ because it has been demonstrated that the myogenic mechanism is typically dominant in this regulatory process^44,49^. The myogenic response is driven by changes in transmural pressure *p*_*t,ij*_, relative to its baseline value *p*_*to,ij*_. The endothelial response depends on changes in wall shear stress *τ*_*ij*_, compared to its baseline value *τ*_*o,ij*_. The baseline values, *p*_*to,ij*_ and *τ*_*o,ij*_, correspond to the values obtained from the baseline flow and pressure fields. Additionally, aligned with *in vivo* studies^66–68^, a threshold of *τ*_*ij*_ =0.1Pa was adopted to ensure that only physiologically relevant levels of shear stress elicit a regulatory response. The wall shear stress, *τ*_*ij*_, in each vessel was estimated assuming a parabolic velocity profile typical for laminar pipe flow:

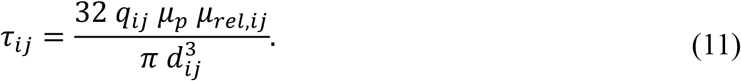

As compliance can only vary within set limits from its baseline value, we applied a sigmoidal function to link the stiffness *k*_*ij*_ to relative compliance *C*_*r,ij*_ ^40,42,49^:

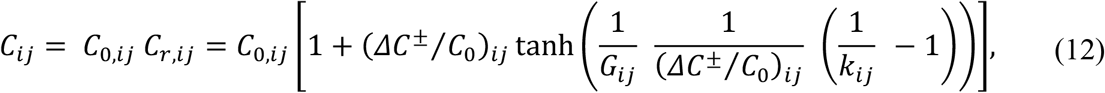

where (Δ*C*^±^/*C*_0_)_*ij*_ defines the bounds for positive and negative compliance changes (asymmetrically constrained) and *G*_*ij*_ sets the steepness of the sigmoid slope. The values of parameters (*S*_*σ,ij*_, *S*_*τ,ij*_, *G*_*ij*_,(Δ*C*^±^/*C*_0_)_*ij*_) were selected based on values reported in prior computational studies^49^ (see Table S1).

By computing the compliance according to Eqs. (10) and (12) and subsequently updating parameter *β*_*ij*_ according to Eq. (8), changes in compliance directly modify the pressure-area relationship and allow the vascular system to adapt to pressure changes and to reach a new equilibrium. By substituting Eq. (3), (8), and (12), the diameter *d*_*ij*_ of regulatory vessels under active control can be directly computed as:

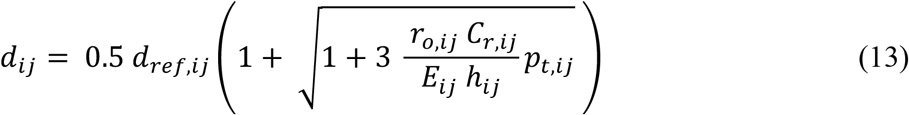

By looking at Eqs. (6) and (13), which describe how vessel diameters are updated under passive and active mechanisms, we observe that the minimum achievable diameter corresponds to the reference diameter *d*_*ref*_, occurring when the transmural pressure *p*_*t*_ approaches zero. This lower bound is an inherent constraint of our model and was further tested and justified based on *in vivo* experimental observations^5^. As pressure and diameter are mutually dependent, the system needs to be solved iteratively. At each iteration, pressure and flow rate distributions were first computed with the blood flow model (Eqs. (1) and (2)). Using the updated pressures and flow rates, for the regulatory vessels, we evaluated the transmural pressure and shear stress to determine the relative stiffness (Eq. (10)). These values were subsequently used to update vascular compliance (Eq. (12)) and compute new vessel diameters (Eqs. (13)). For the passive vessels, diameter changes were solely driven by pressure variations, according to Eq. (6). The loop was repeated until convergence, defined as a maximum relative change in diameter below 10^-6^, was reached. This was typically achieved within 150 iterations. A schematic representation of our autoregulatory model is illustrated in Figure 1B. All parameter values used in our *in silico* model are provided in Table S1.

A sensitivity analysis of key input parameters governing the autoregulatory response, including the myogenic and endothelial sensitivity factors (*S*_*σ,ij*_, *S*_*τ,ij*_), the slope parameter (*G*_*ij*_), and the bounds on compliance variation (Δ*C*^±^/*C*_0_)_*ij*_ has been performed and is summarized in Appendix A. By varying these parameters within ±25% of their respective baseline values, we observed that the resulting changes in model output remained below 10%, confirming the robustness of the model’s autoregulatory behavior.

## Results

Our results focus on analyzing cerebral autoregulatory behavior for different physiological and pathological scenarios, namely (i) healthy, (ii) stroke-reperfusion, and (iii) chronic autoregulation dysfunction conditions, as shown in Figure 1C. Each setup was defined to investigate specific aspects of autoregulatory control and vascular adaptation within our microvascular networks. In the following, the specific simulation setup and the results for the different scenarios are presented jointly to facilitate the interpretation of the results.

### Vascular Responses Are Heterogeneous and Vary with Arterial Location

To mimic autoregulation in healthy conditions, we started from the baseline state of our four networks and varied the MAP (inflow pressure) within the range of 50–140 mmHg, while maintaining active diameter regulation in all arterial segments (see also Figure 1B). To validate the static autoregulation model, we reconstructed the classic static autoregulation curve, which shows how CBF responds to changes in MAP (Figure 2A). The modelled static autoregulation curves align well with *in vivo* data^69^, confirming the validity of our *in silico* model. For further analysis, we defined the autoregulatory range as the MAP interval where CBF remains within ±10% of its baseline value. Based on the average curve, this range was identified as approximately 68 – 118 mmHg, consistent with experimental observations^15,69^.

**Figure 2.**
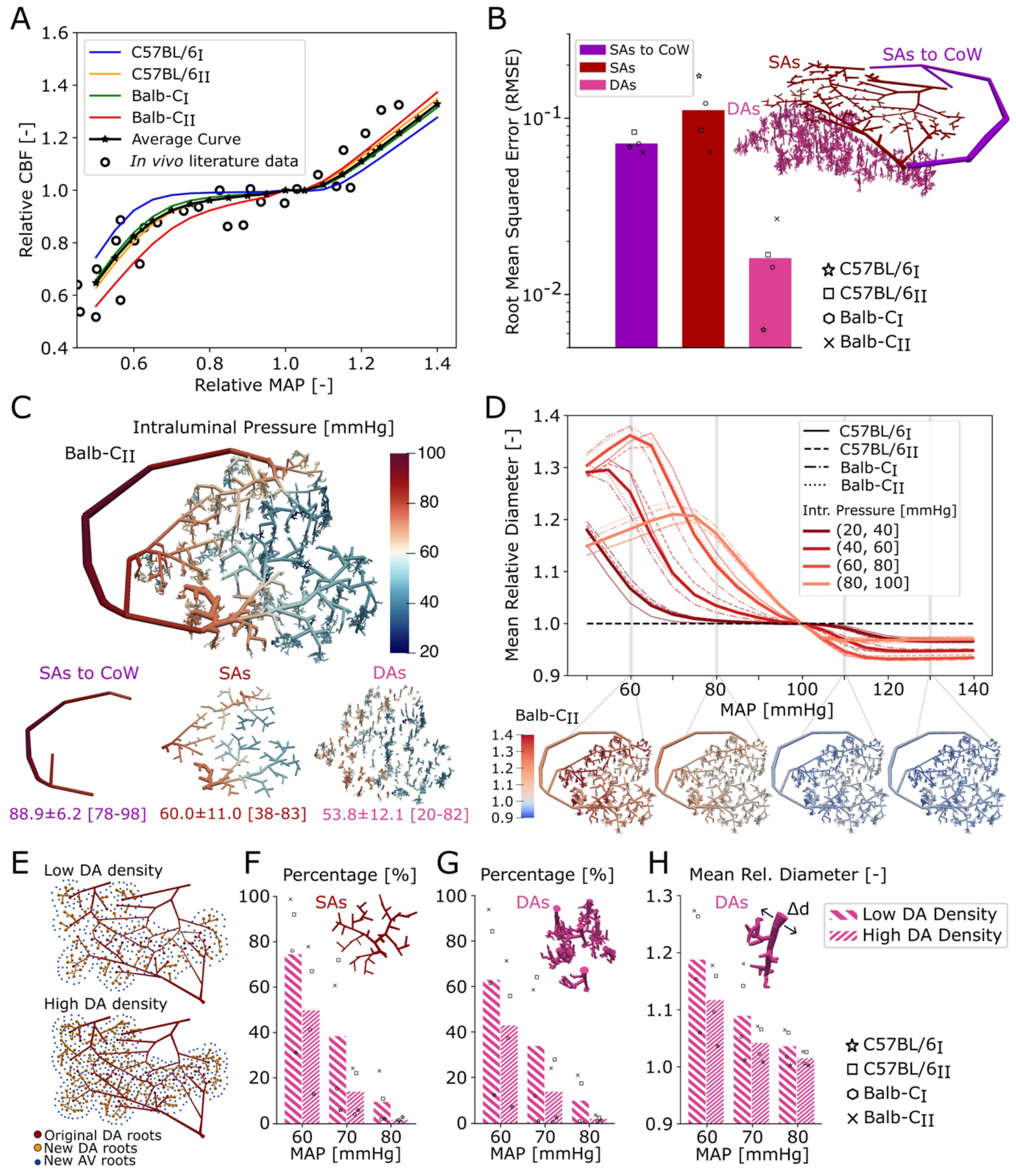
The role of different arteriole levels for cerebral autoregulation under healthy conditions. **(A)** Static autoregulation curves for all microvascular networks compared to *in vivo* experimental measurements^69^. **(B)** Root Mean Squared Error (RMSE) between the fully regulated and the modified autoregulation curve. The modified curves for each network are computed by deactivating different arterial groups, namely: (i) surface arteries connecting to the circle of Willis (SAs-to-CoW), (ii) surface arteries (SAs), and (iii) descending arterioles (DAs). **(C)** Intraluminal pressure distribution across all arterial segments in the Balb-CII microvascular network at baseline conditions. Summary statistics (mean ± SD and range) are reported for each arterial group. **(D)** Mean relative diameter change as a function of MAP across four baseline intraluminal pressure intervals: (20, 40], (40, 60], (60, 80], and (80, 100] mmHg. The non-solid lines show the averaged relative diameter change across all vessels within each pressure group and the solid line the average across networks. The bottom panel shows spatial maps of relative diameter across the Balb-CII network at selected MAP values, highlighting how vasodilation and vasoconstriction evolve with pressure. **(E)** Two surface SA topologies derived from C57BL/6I mouse illustrating low and high DA root densities, respectively. **(F-G)** Proportion of vessels showing >10% dilation in response to MAP drops (60, 70, 80 mmHg) across different DA densities: **(F)** SAs, and **(G)** DA trees (mean dilation per tree). **(H)** Mean relative diameter across all DA trees at 60, 70, and 80 mmHg, comparing low and high DA density cases. For each DA tree, the mean relative diameter was computed, followed by averaging across all trees in the network.

In the subsequent analysis, we focused on the contribution of different arterial segments to overall autoregulation. To quantify the role of different arterial groups for maintaining CBF constant across the autoregulatory range, we deactivated the autoregulatory ability in three arterial groups independently, namely: (i) SAs-to-CoW), (ii) SAs, and (iii) DAs (Figure 2B). In each case, the corresponding arteries responded passively to MAP changes, and a modified autoregulation curve was calculated. The root mean squared error (RMSE) relative to the fully regulated (control) curve is smallest for passive DAs, suggesting that SAs-to-CoW and SAs are able to compensate for the lacking active DA vasodynamics (Figure 2B). This is not the case for deactivated vasodynamics at the level of SAs-to-CoW and SAs, highlighting their prominent role for healthy autoregulation. Figure S1 shows the mean autoregulation curves across all networks for each scenario.

To further investigate how individual arteries contribute to MAP changes, we spatially quantified the relative diameter changes in terms of intraluminal pressure (IP), which serves as a proxy for the arterial location within the network (Figure 2C). More precisely, arteries with higher IP are generally found in vessels located closer to the CoW. The most upstream arteries (IP: 80-100 mmHg) responded to smaller MAP variations, reaching maximum dilation at a MAP of approximately 75 mmHg. In contrast, more distal arteries (IP: 60-80 mmHg) only reacted when the pressure drop became more pronounced, reaching their peak dilation at around 60 mmHg and exceeding the dilation observed in the most upstream vessels (Figure S2A). Interestingly, these upstream arteries tend to exhibit smaller dilation amplitudes and reach their maximal response at relatively modest pressure drops, while smaller, downstream arteries display a greater capacity to dilate under larger MAP reductions. Regarding vasoconstriction, our model predicts that arterial diameters decrease by less than 10% relative to baseline in response to elevated MAP levels, and this constricted state is sustained even with further increases in MAP. This behavior reflects the lower diameter bound imposed by the reference diameter *d*_*ref*_ (see Discussion for further justification based on *in vivo* evidence). More specifically, arteries located closer to the CoW (IP: 90-100 mmHg) appear to exhibit their strongest constriction following a 10% increase in MAP and remain constricted with subsequent MAP elevations. Arteries with baseline intraluminal pressures between 70–80 mmHg showed the largest diameter reductions across all levels of MAP increase (Figure S2B).

### High DA density increases vascular autoregulatory capacity

As noted previously, our microvascular networks differ in their density of DAs; therefore, we quantified how this anatomical feature influences the spatial distribution of the autoregulatory response. To this end, we artificially adjusted the DA density in our four original networks (2× C57BL/6 and 2× Balb-C) to generate four networks with high DA density and four with low DA density, matching values reported in the literature for C57BL/6 (13.4 DAs/mm^2^) and Balb-C (8.9 DAs/mm^2^), respectively^51^(Figure 2E). Importantly, LMC characteristics were maintained, and only DA root density was modified (step b, in Figure 1A).

Although the overall shape of the autoregulation curve remained relatively consistent across networks with different DA densities, we observed more extensive arterial dilation in the networks with lower DA density (Figure S2). Networks with fewer DAs exhibited a higher proportion of SAs and DA trees dilating by at least 10% in response to a MAP decrease (Figure 2F, G). For example, in networks with fewer DAs, twice as many SAs dilated by at least 10% following a 30% MAP reduction (70 mmHg) compared to networks with higher DA density (38.5% vs. 14.1%, Figure 2F). Additionally, DAs exhibited more pronounced dilation in these low-density configurations, with a 40% decrease in MAP (60 mmHg) leading to 18.8% dilation, compared to 11.7% in high-density networks (Figure 2H). In summary, this suggests that networks with a low DA density might be more susceptible to autoregulatory impairments, while networks with a high DA root density are characterized by a higher autoregulatory capacity.

### Autoregulatory Failure Amplifies Capillary Flow Instability Across MAP Variations

Cerebral autoregulation can become impaired following ischemic stroke, resulting in unadjusted CBF in response to pressure variations. As vessels on the MCA side, downstream of the previously occluded segment, are most likely to be affected during stroke, we defined two approaches to select arteries that lose autoregulatory capacity: (i) upstream-to-downstream: starting from larger upstream SAs and progressively including smaller downstream arteries, and (ii) downstream-to-upstream: beginning with the most distal SAs and extending the loss of regulation toward more proximal vessels. In both strategies, the DAs extending from the impaired SAs were also assumed to be affected. For each strategy, we quantified the minimum number of impaired arteries required to significantly alter the autoregulation curve and assessed how such impairments affected capillary perfusion. Similarly to healthy conditions, the baseline networks are used for this study, and MAP is varied within the range of 50–140 mmHg.

For both strategies, the extent of autoregulatory impairment was quantified as the RMSE between the original (fully regulated) autoregulation curve and the progressively impaired curve. This RMSE was then normalized by the RMSE between the fully regulated curve and the fully impaired case, in which all arteries on the MCA side downstream of the previously occluded segment were assumed to have lost their autoregulatory function (i.e., passive vessels, Figure S4A). An impairment score of 100%, therefore, indicates that the impaired autoregulation curve is indistinguishable from the fully impaired (passive) case. The original, ~55% impaired and 100% impaired curves are illustrated in Figure S4B.

#### Upstream-to-downstream impairment

All MCA-sided surface arteries (SAs-to-CoW and SAs) downstream of the occluded vessel are divided into generations, where each generation represents a new bifurcation (Figure 3A). Our results showed that impairing arteries up to the third generation resulted in approximately 50% autoregulation impairment, indicating that these arteries play a key role in regulating blood flow in response to pressure variations (Figure 3B). This finding suggests that even when recanalization is achieved, the loss of regulatory function in the first major artery downstream of the occlusion can substantially impair perfusion under varying MAP conditions.

**Figure 3.**
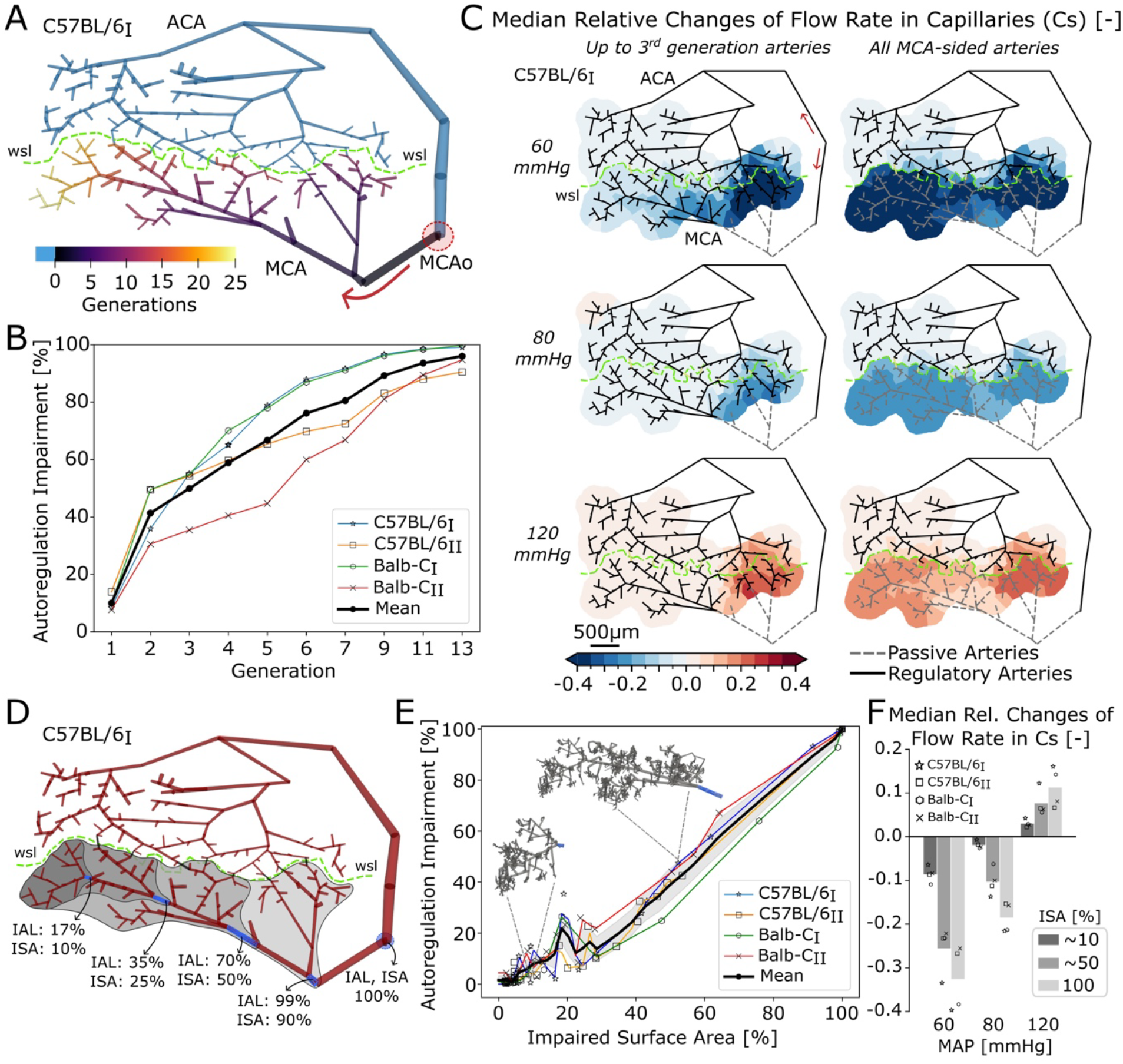
Stroke-induced autoregulatory impairment based on generation- and area-specific analysis. **(A)** Arterial generation map for the C57BL/6I network, used to implement the *upstream-to-downstream* impairment approach. First-generation arteries are defined as those immediately downstream of the previously occluded segment (marked by the red dashed circle), with each subsequent generation defined by successive bifurcations. **(B)** Autoregulation impairment as a function of the number of impaired arterial generations. Results are shown for each individual network, along with the average trend across all networks. **(C)** Spatial maps of median relative changes in capillary flow rate (Cs) at three mean arterial pressure (MAP) levels (60, 80, and 120 mmHg), for two impairment scenarios: (i) impairment restricted to surface arteries up to the 3rd generation (left), and (ii) impairment of all arteries on the MCA side (right). Passive and regulatory arteries are indicated with dashed and solid lines, respectively. The relative change is computed with respect to the fully regulated case at each pressure level. **(D)** Surface topology of the C57BL/6I network showing progressive impairment of surface arteries, used in the *downstream-to-upstream* impairment approach. Impairment is quantified in terms of impaired arterial length (IAL) and impaired surface area (ISA), both expressed as percentages. The location with 100% IAL and ISA corresponds to the site of the original occlusion. **(E)** Autoregulation impairment as a function of impaired surface area. Results are shown for each network and as an averaged curve across all networks. Insets illustrate representative surface artery configurations corresponding to different ISA levels. **(F)** Median relative changes in capillary flow rate (Cs) at 60, 80, and 120 mmHg for three levels of impairment: ~10%, ~50%, and 100% impaired surface area. Cs was selected with 300μm of impaired SAs. The relative change is computed with respect to the fully regulated case at each pressure level. Bars represent mean across four networks.

To evaluate how this impairment spatially affects perfusion at the capillary level, we computed the median relative changes in flow rate for capillary columns positioned below DA root points. We considered three MAP values, *i.e*., 60, 80, and 120 mmHg, and two impairment scenarios: (i) impairment limited to surface arteries up to the third generation, and (ii) complete impairment of all arteries on the MCA side (Figure 3C). At 60 mmHg (40% below baseline), both impairment cases resulted in a marked reduction in capillary flow across the network (Figure 3C, first row). Under full MCA impairment, the median relative flow change in all MCA-sided capillaries reached −42%, whereas the third-generation impairment produced a more localized but still substantial decrease. This highlights the inability of passive vessels to sustain adequate perfusion under severely hypotensive conditions. At 80 mmHg the flow reduction was less pronounced. Nonetheless, for the full MCA impairment, the median relative capillary flow change was −20%, indicating that even moderate pressure drops can lead to significant perfusion deficits when autoregulation is compromised. In contrast, at 120 mmHg, the loss of autoregulation led to capillary hyperperfusion, as passive arteries were unable to buffer the elevated perfusion pressure. This resulted in a widespread increase in flow across the network, with the full MCA impairment condition exhibiting a particularly pronounced effect, characterized by a median relative flow increase of +16% in MCA-sided capillaries.

Overall, these results demonstrate that the spatial impact of autoregulatory failure is modulated by both the magnitude of MAP deviation from baseline and the extent of arterial impairment. While localized impairment leads to moderate region-specific effects, the loss of regulation in all MCA-sided arteries results in widespread perfusion disturbances, with changes propagating across the entire microvascular network. This highlights the crucial role of proximal surface arteries in maintaining microvascular perfusion under both hypotensive and hypertensive conditions.

#### Downstream-to-upstream impairment

In the downstream-to-upstream impairment approach, surface arteries were progressively impaired based on their perfusion levels during stroke, under the assumption that more severely hypoperfused vessels are more susceptible to losing their autoregulatory capacity (Figure S5). Figure 3D illustrates a representative network divided into four spatial regions, each exhibiting a distinct degree of impairment. These regions are characterized by the percentage of impaired artery length (IAL) and impaired surface area (ISA), reflecting the cumulative progression of impairment from distal branches toward proximal arteries.

To quantify the impact of this impairment on autoregulation, we examined the relationship between the percentage of impaired surface area and the resulting impairment in autoregulation, as previously defined (Figure 3E). Our results showed that when approximately 45–50% of the surface area was impaired, the corresponding autoregulatory impairment reached moderate levels of around 40%. This suggests that even partial spatial loss of autoregulatory function, concentrated in distal regions, can result in a measurable decline in the network’s ability to maintain stable perfusion across varying pressure conditions.

To evaluate how this progressive impairment affects microcirculatory perfusion, we calculated the median relative change in capillary flow under three MAP conditions (60, 80, and 120 mmHg) and for three levels of impaired surface area (~10%, ~50%, and 100%). Cs were selected within a planar distance of 300 μm to impaired SAs. At 60 mmHg, the perfusion deficit was most pronounced, with a mean reduction in capillary flow of approximately −34% under full MCA impairment (Figure 3F). At 80 mmHg, the mean flow reduction was around −20%, while at 120 mmHg, the capillary beds exhibited hyperperfusion, with flow increases reaching approximately +12%. These trends closely mirrored those observed under upstream-to-downstream impairment, reinforcing the conclusion that the spatial extent of surface artery dysfunction critically modulates the pressure-dependent flow response at the capillary level. Figure S6 shows spatial maps of median relative changes in capillary flow rate at three MAP levels (60, 80, and 120 mmHg), for the four impairment scenarios as shown in Figure 3D.

Together, these results confirm that persisting vascular impairments following stroke, *i.e*., arteries that lose their ability to autoregulate, lead to impaired autoregulation curves. Importantly, the degree of impairment scales with the severity of stroke-related vascular dysfunction.

### Altered Vasodynamics are a Plausible Cause for CBF Overshoot during Reperfusion

As previously described, *in vivo* experiments have shown that reperfusion dynamics following ischemic stroke may depend on the extent of LMCs^37^. A prevailing hypothesis suggests that poor collateral networks are more prone to perfusion overshoot due to ischemia-induced endothelial damage, which impairs the regulatory capacity in the recanalized MCA. In contrast, rich collateral networks may sustain minimal flow during occlusion, thereby preserving endothelial integrity and enabling more physiologically regulated reperfusion upon recanalization. In this context, we addressed the question whether the absence of LMCs alone could, from a hemodynamic perspective, cause the perfusion overshoot observed experimentally in Balb-C (LMC-poor) mice. To isolate the effect of LMCs, we artificially modified the number of LMCs in each of our four original networks (derived from two C57BL/6 and two Balb-C mice), creating two configurations per network: one with abundant collaterals (100% LMCs) and one without collaterals (0% LMCs). These paired configurations differ only in the number of LMCs and are otherwise identical.

To mimic middle cerebral artery occlusion (MCAo), a 50 μm segment of the MCA-sided SAs-to-CoW (M2 segment) artery was constricted to 10% of its baseline diameter. MAP was kept constant at 100 mmHg and vessel diameters were adjusted in response to the occlusion (*i.e*., stimulus: MCAo). To obtain representative diameter adjustments under stroke conditions and to isolate the role of autoregulatory mechanisms in shaping reperfusion dynamics, we adopted a computational strategy in which all vessels, except LMCs, were assumed to respond passively during stroke and actively during reperfusion. This assumption reflects the ischemic environment, in which perfusion pressures fall below the typical autoregulatory range, resulting in a predominantly passive vascular response. This simplification allowed us to decouple hemodynamic effects caused by reduced perfusion pressure from those arising due to impaired or restored vascular regulation. In contrast, LMCs retained active autoregulation throughout both stroke and reperfusion phases, consistent with experimental evidence showing their vasodilatory behavior during ischemia. Passive behavior during the occluded phase reflects the ischemic environment in which regulatory responses are often suppressed, while the reintroduction of active regulation during reperfusion enables a controlled evaluation of how vessel responsiveness modulates flow restoration. Moreover, *in vivo* studies suggest that pial vessels tend to either reduce their diameter^70^ or show no significant diameter changes^71^ under stroke conditions, further supporting the assumption of passive behavior in the model.

During reperfusion, we simulated tissue plasminogen activator (tPA) treatment by stepwise reopening the occluded vessel (Figure 4A). At this stage, the stroke networks served as the baseline case, *i.e*., all baseline variables – denoted with subscript *o* - have been updated accordingly. MAP is held constant at 100 mmHg throughout the gradual reopening. Vessel diameters now change in response to pressure redistributions caused by the reopening of the occluded segment (*i.e*., stimulus: gradual vessel reopening).

**Figure 4.**
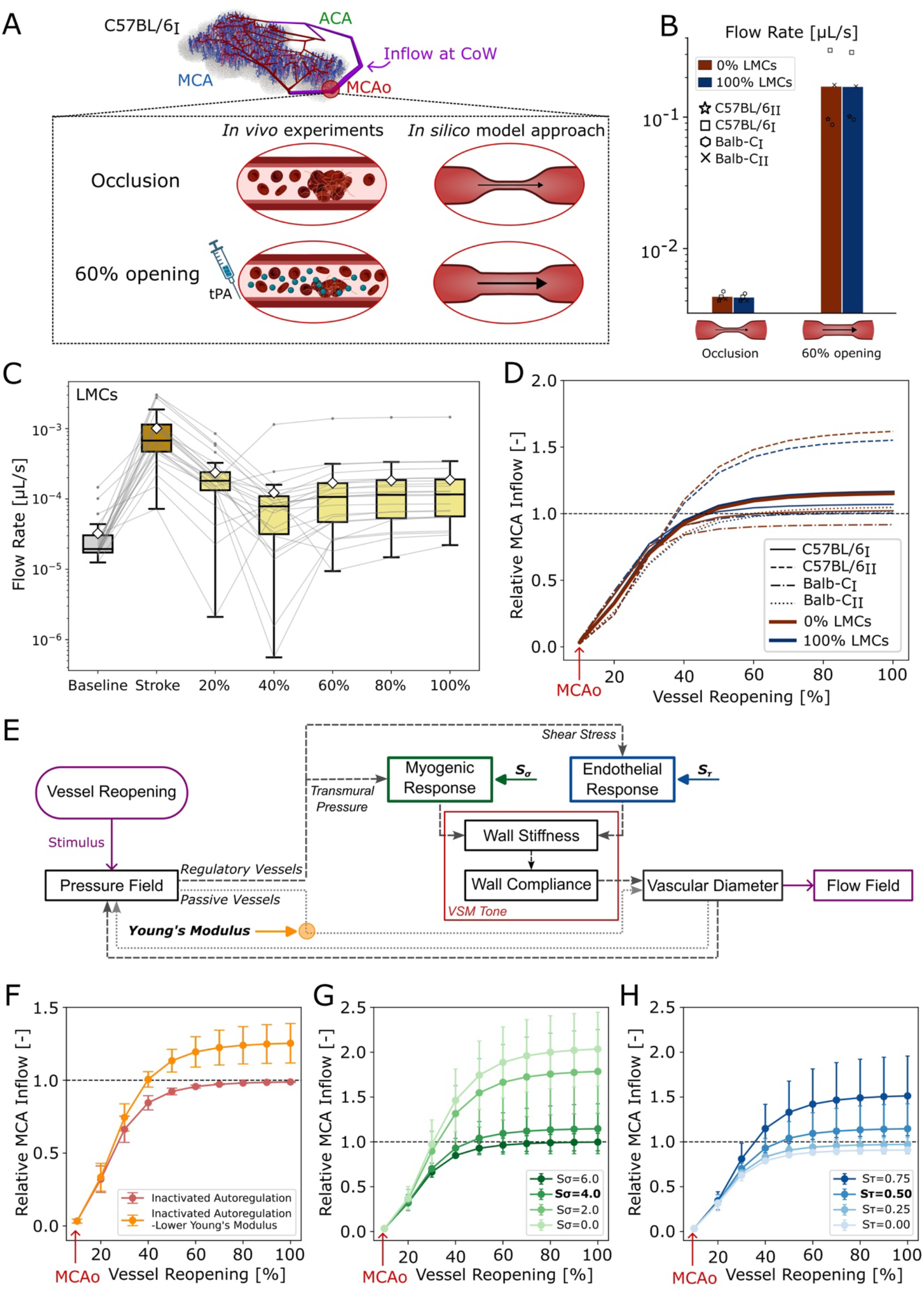
Interaction of vascular mechanisms in reperfusion after MCA occlusion. **(A)** Schematic representation of the C57BL/6I network showing the occlusion site in the MCA and the effect of tPA treatment at the clot location (red circle). **(B)** Flow rate [μL/s] through the occluded vessel during stroke (vessel diameter at 10% of baseline) and reperfusion (vessel reopened to 60% of baseline) conditions, shown for 0% and 100% leptomeningeal collaterals (LMCs). Bars represent mean across all networks. **(C)** Flow rate [μL/s] through the LMCs under baseline, stroke, and after reopening from 20% to 100%. **(CD)** Relative MCA inflow as a function of vessel reopening (from 10% to 100% of baseline diameter), shown for each network with distinct line styles and for both 0% and 100% LMC conditions. Bold lines indicate the average across all networks for each condition. **(E)** Schematic of the autoregulatory model for reperfusion highlighting mechanisms potentially affected by vascular impairments in response to stroke. Independently altered characteristics are myogenic (*S*_*σ*_) and endothelial (*S*_*τ*_) responses in regulatory vessels, and Young’s modulus in passive vessels. **(F-H)** Relative MCA inflow as a function of vessel reopening under altered regulatory mechanisms: **(F)** the absence of active autoregulation (passive behavior), with and without reduced Young’s modulus; **(G)** active autoregulation with varying myogenic sensitivity (*S*_*σ*_ = 0, 2, 4, 6); **(H)** active autoregulation with varying endothelial sensitivity (*S*_*τ*_= 0, 0.25, 0.5, 0.75). All curves show mean ± SD across networks.

In response to MCA occlusion (MCAo), all arteries, except LMCs, underwent passive constriction of less than 10% from their baseline diameter, while LMCs dilated by approximately 23%, consistent with experimental observations^37,70^ (Table S2). To investigate the role of LMCs during MCAo and reperfusion, we assessed perfusion in the occluded vessel for different network configurations (100% and 0% LMCs) under two conditions: (i) *stroke condition*: the vessel remained constricted at 10% of its baseline diameter, and (ii) *reperfusion conditions*: stepwise reopening of the vessel to 100% of its baseline diameter. Our results show no substantial difference in perfusion in the occluded vessel itself between networks with 100% and 0% LMCs for stroke and 60% reopening (Figure 4B). These results indicate that LMC density does not significantly affect perfusion in the recanalized segment, despite its critical role in regions near the collateral vessels during stroke^51^. Moreover, once the occluded vessel is reopened, the pressure gradient between MCA and ACA territories decreases, reducing the hemodynamic relevance of the LMCs and shifting their flow toward baseline distributions (Figure 4C).

Figure 4D illustrates the total MCA inflow relative to baseline values (*i.e*., before stroke) as a function of vessel reopening, under intact autoregulation in all arteries. The average reperfusion profiles for the 100% and 0% LMC conditions were nearly identical. Interestingly, despite identical model parameters across all networks, the C57BL/6II network exhibited perfusion overshoot in both LMC configurations. This suggests that LMC density may not be the primary driver of perfusion overshoot. Instead, overshoot might result from network-specific variations in the balance between myogenic and endothelial regulation. In contrast, total ACA inflow remains unchanged mainly under these conditions (Figure S7A).

#### Regulatory Determinants of Reperfusion Overshoot

To explore the conditions under which perfusion overshoot might occur, we evaluated several regulatory scenarios to mimic post-ischaemic dysfunction (Figure 4E): (i) Vascular elasticity impairment: Arteries were assumed to lose autoregulatory function and behave passively, with the Young’s modulus (E) of larger vessels reduced to reflect compromised vascular integrity. Specifically, we assumed that only the MCA-sided SAs-to-CoW segments and the first two generations of SAs with diameters >40 μm were affected, and their Young’s modulus was reduced to 10% of baseline. (ii) Myogenic impairment: The myogenic response was either entirely removed or modulated by ±50% of its initial value. (iii) Endothelial impairment: Similarly, the endothelial response was either fully removed or modulated by ±50%. In both (ii) and (iii), the modifications were only applied to all arteries on the MCA side. For these experiments, we used the four original networks, having demonstrated that the extent of LMCs does not significantly influence the reperfusion profile.

In networks where all arteries, except LMCs, behaved passively (*i.e*., without autoregulation), MCA inflow returned to baseline as the occluded vessel reopened (Figure 4F). This outcome is expected since no active mechanisms (feedback) were available to account for changes in vessel parameters (see Eq. (6), *d*_*ref*_ was set constant throughout the simulations). However, when the Young’s modulus of largest upstream vessels was reduced to 10% of baseline, thereby making them more flexible, we observed a clear perfusion overshoot, with relative MCA inflow reaching ~1.3 at 60% reopening. This finding underscores that even in the absence of active regulation, changes in vessel mechanical properties can shape reperfusion dynamics and lead to overshoot.

Subsequently, we varied the myogenic sensitivity factor (*S*_*σ*_) while keeping the endothelial sensitivity fixed at *S*_*τ*_ = 0.5, simulating different levels of myogenic responsiveness (Figure 4G). As *S*_*σ*_ decreased from 6 to 0, the magnitude of MCA inflow during reperfusion increased significantly. For the case of complete myogenic loss (*S*_*σ*_ = 0), relative MCA inflow peaked at ~2.1 at 60% reopening, compared to the strongly regulated condition (*S*_*σ*_ = 6), where flow returned to baseline. This clearly indicates that the loss or weakening of myogenic control facilitates hyperperfusion, supporting the hypothesis that myogenic impairment is a key contributor to reperfusion overshoot, likely because arteries lose the ability to constrict in response to elevated pressure.

If the myogenic response was held constant (*S*_*σ*_ = 4), while *S*_*τ*_ was varied from 0.0 (complete loss) to 0.75 (strong endothelial function) we observed an opposite trend (Figure 4H). Increasing *S*_*τ*_ led to a progressive increase in MCA inflow during reperfusion, with relative inflow rising from ~1.0 (*S*_*τ*_ = 0) to ~1.6 (*S*_*τ*_ = 0.75) at 60% reopening. This suggests that, contrary to the myogenic response, a strong endothelial response may amplify reperfusion overshoot, likely due to excessive vasodilation in response to elevated shear stress. Thus, endothelial upregulation may contribute to post-stroke hyperemia.

Together, our findings demonstrate that impaired autoregulation, particularly myogenic dysfunction, could be a primary driver of reperfusion overshoot. A reduced Young’s modulus and elevated endothelial sensitivity can further amplify the response. However, loss of myogenic tone yields the most pronounced overshoot among the tested conditions. LMCs on the other hand are mostly relevant during MCAo and likely by avoiding vascular damage at this stage. These results highlight the intricate interplay between passive mechanical properties and regulatory mechanisms in shaping cerebral hemodynamics after stroke and may provide insight into why some patients experience excessive perfusion during reperfusion therapy.

## Discussion

Using a novel computational framework that incorporates the two principal autoregulatory mechanisms, namely myogenic and endothelial regulation, we present unprecedented insights into how the cerebral microvasculature responds under (i) healthy autoregulation, (ii) ischaemic stroke–reperfusion with altered vascular reactivity, and (iii) chronic autoregulatory dysfunction after stroke.

In the context of healthy autoregulation, we observed that the autoregulatory response is not uniform but depends strongly on an artery’s location and baseline intraluminal pressure. Larger surface arteries extending from the CoW were the primary responders to moderate MAP changes, whereas downstream arteries showed smaller diameter changes. Under more pronounced hypotension, downstream arteries exhibited marked vasodilation, indicating a secondary but significant contribution to flow regulation under more extreme conditions. These findings align with *in vivo* studies^4,5,17^ and underscore the importance of proximal vessels in pressure regulation. Interestingly, strain-dependent variability in DA density modulated this response. Networks with lower DA density required larger diameter changes in surface arteries to maintain perfusion, leading to stronger and more spatially widespread responses. This observation puts forward that a high DA density is beneficial for increase autoregulatory capacity and might have a protective effect towards autoregulatory impairments during stroke. While differences in DA density have been reported for different mouse strains^52^, it remains unknown if DA density might also vary between human individuals and if this could be contributing to differences in stroke outcome.

Moreover, our results show that autoregulatory failure in the territory most affected during stroke can significantly alter the cerebral autoregulation curve. For instance, even a modest 20% drop in MAP, such as may occur during sleep^72,73^, can lead to a 16% reduction in total CBF when all MCA-sided arteries are impaired, compared to only a 4% reduction in networks with intact autoregulation. A persistent hypoperfusion during sleep could reduce the brain’s important ability to clear metabolic wastes, leading to the accumulation of tau tangles/amyloid-β^74–76^ – a known precursor during dementia^77,78^. These findings suggest that local autoregulatory failure after stroke may increase the brain’s vulnerability to neurodegenerative processes^79–81^ during otherwise benign physiological states, such as sleep.

Our simulation framework allows us to precisely quantify the impact of a global CBF reduction for individual capillary perfusion - a metric very challenging to assess in *in vivo* experiments, however, highly relevant for local oxygen and nutrient supply. In this context, our results confirm that impaired regulation in surface arteries propagates throughout the microvascular network and consequently plays an important role in a well-regulated oxygen and nutrient supply.

By testing two complementary approaches for autoregulatory impairment — progressively increasing the number of affected vessels either from the upstream to the downstream side or in the reverse direction — we uniquely implement a stepwise, spatially controlled approach to assess the impact of partial regulation loss. Our findings provide further evidence that proximal surface arteries play a central role in maintaining stable microvascular perfusion under both hypotensive and hypertensive conditions. However, even the loss of autoregulation at a few distal artery branches causes notable local hypo- or hyperperfusion at the capillary level. Importantly, the spatial impact of regulatory failure on capillary perfusion depends on both the magnitude of the MAP deviation and the extent of arterial impairment. These results may also have implications for neurovascular coupling (NVC). Since both autoregulation and NVC depend on the healthy function of VSMCs and endothelial cells, ischemic injury that compromises these components may simultaneously impair both mechanisms. In this context, the stroke-induced regional loss of vessel responsiveness may not only impair pressure-driven flow regulation but also reduce the brain’s capacity to match blood supply to neuronal activity^77,82–85^.

Exploiting the benefits of our *in silico* framework and generating networks that only differ in LMC density, we showed that LMC extent had limited influence on perfusion in proximal arteries, particularly within the recanalized MCA. This suggests that perfusion in the most upstream regions may be largely independent of the extent of LMCs. Consequently, based on our simulations, the hypothesis that LMCs contribute to maintaining sufficient perfusion to the recanalized MCA and thereby enable it to remain normally regulated may not be fully valid. However, LMC extent clearly contributes to elevating perfusion levels during stroke^51,86–88^, which likely helps to protect vessels from autoregulatory impairments^86^, which, as shown within the current study, is crucial for healthy cerebral autoregulation across the microvasculature.

To better understand the conditions leading to hyperperfusion during reperfusion, we examined several scenarios involving variations in vessel wall stiffness and autoregulatory function. Our simulations showed that reperfusion overshoot can occur even in the absence of active regulation, especially when arteries become more flexible (*e.g*., lower Young’s modulus values). However, the most pronounced overshoot was observed when the myogenic response was impaired, emphasizing the importance of vasoconstrictive control in preventing excessive perfusion after recanalization. This finding aligns with experimental studies that point to mechanisms such as peroxynitrite formation and oxidative stress as contributors to diminished vascular tone and myogenic reactivity during reperfusion. Specifically, work by Cipolla *et al*.^13,14,89^ demonstrated that the duration of ischemia and reperfusion (I/R) strongly determines the loss of MCA myogenic activity and tone, with more prolonged I/R producing greater impairment. More recently, Coucha *et al*.^90^ showed that I/R-generated peroxynitrite and protein nitration reduce myogenic tone in MCAs from both the ischemic and contralateral hemispheres, implicating oxidative/nitrative modification as a bilateral mechanism of myogenic dysfunction. These experimental observations have motivated the suggestion that preserving or restoring myogenic responsiveness could be a therapeutic strategy to limit reperfusion-associated hyperperfusion and downstream injury. Taken together, our findings suggest that variability in reperfusion dynamics is likely driven by differences in autoregulatory responses and impairments across mouse strains rather than solely by the extent of collaterals.

Importantly, impaired vascular tone is not unique to stroke. A growing body of evidence shows that myogenic dysfunction also occurs in other conditions, such as aging, hypertension, diabetes, and Alzheimer’s disease, where it contributes to cerebrovascular dysregulation and increased vulnerability to flow disturbances^82,91,92^. Thus, our results may be broadly relevant to understanding flow abnormalities in diverse pathologies characterized by loss of vascular reactivity, beyond the context of ischemic stroke alone.

While our observations for healthy autoregulation generally align well with experimental studies of static autoregulation, the maximum achievable constriction in our simulations is ~10%. This is a consequence of the definition of the reference diameter, which sets a limit for the smallest attainable diameter. Although previous *in vivo* and computational studies have reported similar maximum levels of vasoconstriction across a broad MAP range^5,9,41,93,94^, some vascular segments may exhibit stronger responses under extreme hypertensive conditions, causing an underestimation of vasoconstrictive responses at the single vessel level. However, for most scenarios, this will be compensated by a larger fraction of vessels attaining the largest level of constriction. Only for extreme hypertensive states, our simulations may underestimate perfusion changes and not fully capture the accurate redistribution of flow at single vessel resolution.

Additionally, while we compared networks based on C57BL/6 and Balb-C mouse strains, the autoregulatory parameters might differ between strains and were not tailored to strain-specific behavior. In theory, strain-specific vascular properties and regulatory sensitivities could be easily incorporated in the model to better capture biological variability. This may be particularly relevant given existing evidence of strain-dependent inflammatory responses and vascular remodeling^95,96^. However, precise *in vivo* quantification of such parameters is still lacking and thus needs to be addressed in future studies. Importantly, we are confident that such refinements wouldn’t affect our overall conclusions on the role of different arterial segments for healthy autoregulation and the consequences for impaired autoregulation on healthy brain function.

To the best of our knowledge, this is the first *in silico* study to incorporate static autoregulatory mechanisms to investigate how individual arteries contribute to cerebral blood flow regulation, and how the failure of these mechanisms alters perfusion across the entire microvasculature. Our simulations quantitatively demonstrate the dominant role of surface arteries in buffering MAP changes. Networks with lower penetrating arteriole density exhibited more extensive dilations, reflecting the influence of vascular topology on diameter changes during autoregulation. Impairment of proximal surface arteries led to the most substantial disruptions in microvascular perfusion in response to pressure variations. We also show that altered vasodynamics, such as changes in vessel stiffness or regulatory response, can plausibly cause CBF overshoot during reperfusion.

Given that reperfusion injury and failure remain frequent and clinically relevant complications in stroke treatment, there is a strong need to explore new therapeutic strategies. Our simulations suggest that preserving or restoring myogenic tone may serve as a promising therapeutic target to prevent hyperperfusion, reduce the risk of hemorrhagic complications, and improve outcomes following recanalization^89,90^. These findings highlight the importance of assessing whether vasculature has recovered regulatory function after acute treatment, as persistent vasomotor dysfunction may lead to long-term complications such as impaired perfusion control.

## Acknowledgments

We thank Dr. med. Hakim Baazaoui for his support and valuable discussions.

## Appendix A Sensitivity Analysis of Autoregulatory Model Parameters

Precise values for input parameters in such models often lack direct experimental validation. Hence, given the uncertainty in the key input parameters governing our autoregulation model, we performed a parameter sensitivity analysis to quantify the relative influence of each input on the model output. Specifically, we examined the sensitivity of our model to four critical parameters: the myogenic sensitivity factor (*S*_*σ*_), the endothelial (shear-dependent) sensitivity factor (*S*_*τ*_), the slope of the sigmoidal compliance-stiffness relationship (*G*), and the bounds on compliance variation – maximum positive and negative changes in compliance (Δ*C*^±^/*C*_0_).

We conducted this analysis using the Elementary Effects (EE) method, as originally proposed by Morris^1,2^, which enables efficient screening of input factors over a wide parameter space. The EE method has been widely used in computational modeling studies^3^ due to its ability to evaluate the effect of each parameter independently, while requiring significantly fewer model evaluations compared to variance-based methods. Here, the input value ranges were defined as ±25% of their respective baseline values, and model outputs were evaluated at three representative levels of mean arterial pressure (MAP): 60, 80, and 120 mmHg. As model outputs, we considered (i) the relative cerebral blood flow (CBF), and (ii) the mean relative diameter changes in three arterial subgroups: surface arteries connected to the Circle of Willis (SAs to CoW), surface arteries (SAs), and descending arteries (DAs). This framework enabled us to systematically identify which parameters most significantly influence vascular behavior under varying perfusion pressures. C57BL/6I and Balb-CII networks were used.

The analysis yielded three sensitivity metrics: the mean (*μ*_*EE*_), the standard deviation (*σ*_*EE*_), and the mean of the absolute value of the EEs (*μ*_|*EE*|_). The *μ*_*EE*_ metric assesses the overall influence of each factor on the output, and the *σ*_*EE*_ metric measures the variability of these effects, with high *σ*_*EE*_ values indicating non-linear effects or interactions with other parameters.

Figure A1A illustrates the autoregulation curve for both networks with 95% confidence intervals at each tested MAP value. Figure A1B shows the *μ*_|*EE*|_ metric versus *σ*_*EE*_ metric for the relative CBF. At 80 mmHg, *S*_*σ*_ factor emerges as the most influential factor in both networks as indicated by relatively high *μ*_|*EE*|_ and *σ*_*EE*_ values. *G* parameter is the second important factor at this MAP level. However, at the autoregulatory limits (60 and 120 mmHg), Δ*C*^±^/*C*_0_ factor becomes dominant. This shift likely reflects the transition of vessel compliance into saturation regions of the compliance–stiffness curve at extreme MAP values, while at 80 mmHg, compliance remains in the more linear regime, thus diminishing the impact of Δ*C*^±^/*C*_0_.

We demonstrated the relative CBF as a function of *S*_*σ*_ and Δ*C*^±^/*C*_0_ factors to explicitly visualize how the perfusion responds to variations in the most influential parameters (Figure A1C). Within the autoregulatory region (80 mmHg), variations in *S*_*σ*_ have a substantial effect, where higher values lead to greater arterial dilation and restoration of CBF toward baseline levels. In contrast, at the autoregulatory limits (60 and 120 mmHg), changes in *S*_*σ*_ have minimal impact without creating any trends. Meanwhile, Δ*C*^±^/*C*_0_ does not strongly affect CBF at 80 mmHg, but becomes increasingly important at the extreme MAP values. Specifically, higher compliance bounds at 60 mmHg allow for greater vasodilation, while lower compliance bounds at 120 mmHg limit vasoconstriction, both of which lead to elevated CBF. Another noteworthy observation is that, despite varying all input parameters within ±25% of their baseline values, the Balb-CII network did not reach baseline CBF at 80 mmHg for any combination of input values, whereas the C57BL/6I network was able to restore flow closer to baseline under the same conditions. This finding highlights that network topology can also contribute to variability in the model output. Table A1 summarizes the maximum and minimum relative CBF values obtained across the tested parameter space for the two networks, as well as the relative differences between these extrema at each MAP level, calculated as, (*CBF*_*max*_ − *CBF*_*min*_)/ *CBF*_*min*_ × 100).

**Figure A1.**
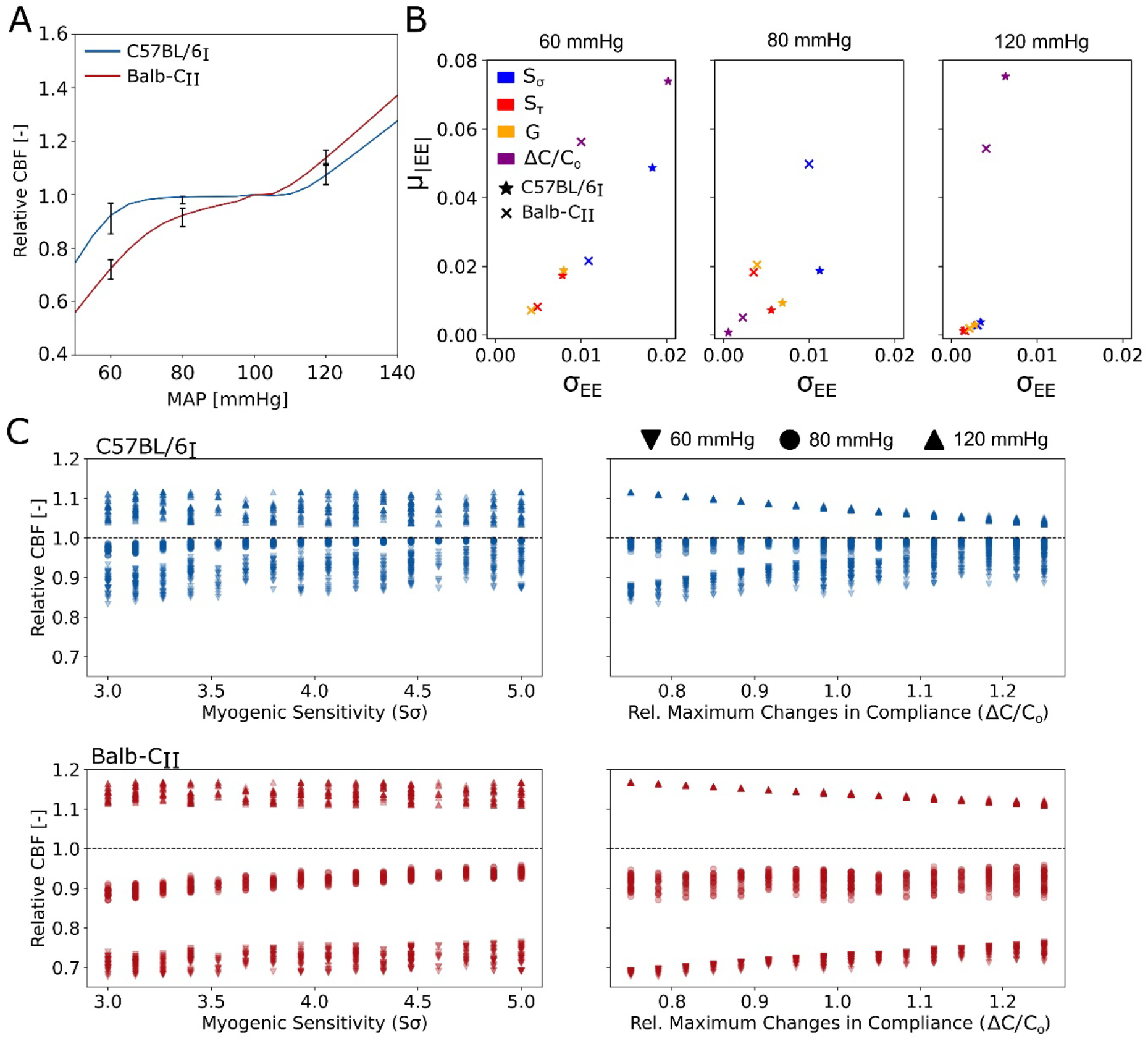
**(A)** Relative cerebral blood flow (CBF) as a function of mean arterial pressure (MAP), shown with 95% confidence intervals based on variations in input parameters at selected MAP levels (60, 80, and 120 mmHg). **(B)** Elementary effects (EE) sensitivity plots showing the mean absolute EE (*μ*|*EE*|) versus the standard deviation of EE (sEE) for relative CBF at different MAP values for C57BL/6I and Balb-CII networks. Parameters analysed include the myogenic sensitivity factor (*S*_*σ*_), the endothelial sensitivity factor (*S*_*τ*_), the slope of the sigmoidal compliance-stiffness relationship (*G*), and the bounds on compliance variation (Δ*C*^±^/*C*_0_). **(C)** Scatter plots illustrating the dependence of relative CBF on changes in myogenic sensitivity factor (*S*_*σ*_) and relative bounds of compliance variations (Δ*C*^±^/*C*_0_) for the two networks at the three different MAP levels. Each data point represents an individual simulation run with parameter values varied ±25% around their baseline values.

**Table A1.** Summary of sensitivity analysis for relative cerebral blood flow (CBF). The table lists the maximum and minimum relative CBF values across the tested parameter space for each mean arterial pressure (MAP) level (60, 80, and 120 mmHg) and for each microvascular network (C57BL/6I and Balb-CII). The percentage (%) column represents the relative difference between these extreme values.

|  | Max value | Min value | Relative Change [%] |
| --- | --- | --- | --- |
| C57BL/6<sub>I</sub> |  |  |  |
| 60 mmHg | 0.98 | 0.83 | 17.9 |
| 80 mmHg | 0.99 | 0.95 | 4.1 |
| 120 mmHg | 1.12 | 1.03 | 7.9 |
| Balb-C<sub>II</sub> |  |  |  |
| 60 mmHg | 0.76 | 0.68 | 12.9 |
| 80 mmHg | 0.96 | 0.87 | 10.2 |
| 120 mmHg | 1.16 | 1.10 | 5.3 |

Figure A2 shows the *μ*_|*EE*|_ and *σ*_*EE*_ matrices for the mean relative diameter changes across three arterial types: surface arteries connected to the Circle of Willis (SAs to CoW), surface arteries (SAs), and descending arterioles (DAs). The trends observed for SAs to CoW vessels closely mirror those seen in the relative CBF analysis, meaning that *S*_*σ*_ is dominant at 80 mmHg, while Δ*C*^±^/*C*_0_ is more influential at the autoregulation limits. However, in the case of SAs and DAs, no single dominant factor is evident at 80 mmHg. Interestingly, at 60 mmHg, while the trends for SAs and DAs in the C57BL/6I network remain consistent with those observed in the SAs to CoW group, the BALB/cII network exhibits a different pattern, indicating that diameter changes in these arterial groups are primarily driven by *S*_*σ*_, followed by Δ*C*^±^/*C*_0_. Additionally, at 120 mmHg, the influence of all parameters appears to diminish across all arterial types. Tables A2 and A3 summarize the relative differences between the maximum and minimum mean relative diameters, *i.e*., (*mean*(*d*)_*max*_ − *mean*(*d*)_*min*_)/ *mean*(*d*)_*min*_× 100), across the parameter space for three arterial types at different MAP levels. Results are shown separately for the C57BL/6I and Balb-CII microvascular networks.

The sensitivity analysis not only clarifies how model outputs are affected by variations in input parameter values, but also offers meaningful insight into the functioning of the autoregulatory response. Our results indicate that the myogenic sensitivity factor, *S*_*σ*_, and the bounds on compliance variation, Δ*C*^±^/*C*_0_, are critical determinants of model behavior across different MAP levels, whereas the endothelial sensitivity factor, *S*_*τ*_, and the slope of the compliance–stiffness relationship, *G*, generally have a more negligible influence on the outcome. Furthermore, we observed that differences in network topology can influence autoregulatory behavior and final outcomes to a notable extent.

**Figure A2.**
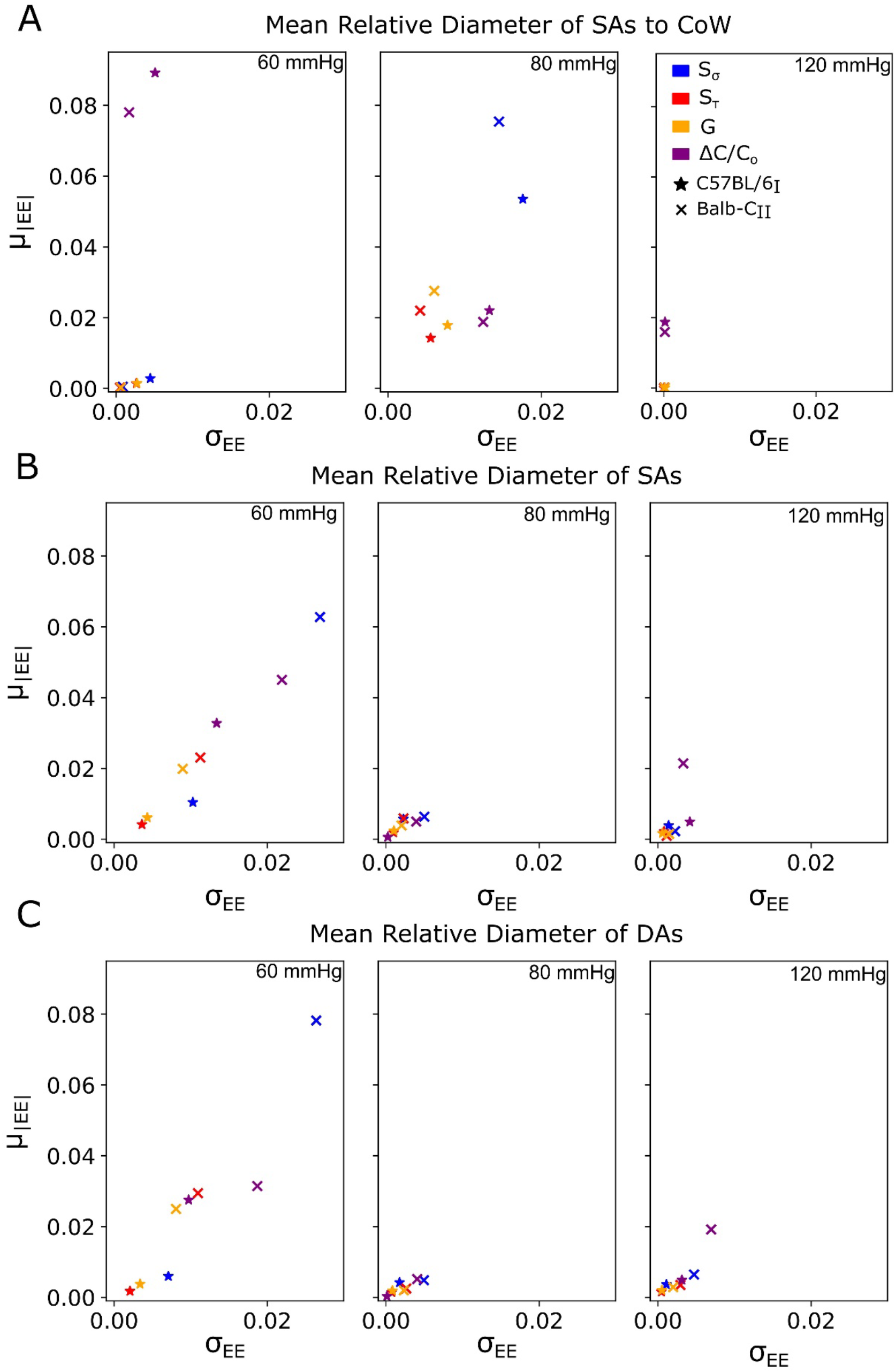
Sensitivity analysis results for mean relative diameters across three arterial types: **(A)** surface arteries connected to the Circle of Willis (SAs to CoW), **(B)** surface arteries (SAs), and **(C)** descending arterioles (DAs). Scatter plots illustrate the mean of the absolute value of elementary effects (*μ*|*EE*|) versus their standard deviation (sEE) at three MAP levels (60, 80 and 120 mmHg) for the C57BL/6I and Balb-CII networks. Each plot identifies the impact of the four tested parameters (myogenic sensitivity factor, *S*_*σ*_; the endothelial sensitivity factor, *S*_*τ*_; the slope of the sigmoidal compliance-stiffness relationship, *G*; and the bounds on compliance variation, Δ*C*^±^/*C*_0_) on diameter responses in the respective arterial group.

**Table A2.** Maximum and minimum mean relative diameters and corresponding relative changes (%) for different arterial groups (SAs to CoW, SAs, and DAs) in the C57BL/6I network at three MAP levels (60, 80, and 120 mmHg). Values reflect the range of model outputs across the tested parameter space in the sensitivity analysis.

|  | Max value | Min value | Relative Change [%] |
| --- | --- | --- | --- |
| 60 mmHg |  |  |  |
| SAs to CoW | 1.23 | 1.13 | 8.10 |
| SAs | 1.10 | 1.04 | 6.18 |
| DAs | 1.07 | 1.01 | 4.84 |
| 80 mmHg |  |  |  |
| SAs to CoW | 1.20 | 1.10 | 9.22 |
| SAs | 1.01 | 1.00 | 0.85 |
| DAs | 1.07 | 1.00 | 0.67 |
| 120 mmHg |  |  |  |
| SAs to CoW | 0.98 | 0.96 | 1.96 |
| SAs | 0.99 | 0.97 | 1.08 |
| DAs | 0.99 | 0.98 | 1.00 |

**Table A3.** Maximum and minimum mean relative diameters and corresponding relative changes (%) for different arterial groups (SAs to CoW, SAs, and DAs) in the Balb-CII network at three MAP levels (60, 80, and 120 mmHg). Values reflect the range of model outputs across the tested parameter space in the sensitivity analysis.

|  | Max value | Min value | Relative Change [%] |
| --- | --- | --- | --- |
| 60 mmHg |  |  |  |
| SAs to CoW | 1.20 | 1.12 | 7.02 |
| SAs | 1.30 | 1.16 | 12.32 |
| DAs | 1.33 | 1.17 | 13.14 |
| 80 mmHg |  |  |  |
| SAs to CoW | 1.23 | 1.09 | 13.03 |
| SAs | 1.08 | 1.06 | 2.84 |
| DAs | 1.07 | 1.05 | 2.06 |
| 120 mmHg |  |  |  |
| SAs-to-CoW | 0.98 | 0.96 | 1.65 |
| SAs | 0.97 | 0.94 | 2.67 |
| DAs | 0.96 | 0.93 | 3.19 |

## References

1. Wang, Y. & Payne, S. J. Static autoregulation in humans. J Cereb Blood Flow Metab 44, 1191–1207 (2024).

2. Silverman, A. & Petersen, N. H. Physiology, Cerebral Autoregulation. in StatPearls (StatPearls Publishing, Treasure Island (FL), 2025).

3. Davis, M. J. & Hill, M. A. Signaling Mechanisms Underlying the Vascular Myogenic Response. Physiological Reviews 79, 387–423 (1999).

4. Kontos, H. A. et al. Responses of cerebral arteries and arterioles to acute hypotension and hypertension. American Journal of Physiology-Heart and Circulatory Physiology 234, H371–H383 (1978).

5. Harper, S. L., Bohlen, H. G. & Rubin, M. J. Arterial and microvascular contributions to cerebral cortical autoregulation in rats. American Journal of Physiology-Heart and Circulatory Physiology 246, H17–H24 (1984).

6. Welsh, D. G. & Segal, S. S. Endothelial and smooth muscle cell conduction in arterioles controlling blood flow. American Journal of Physiology-Heart and Circulatory Physiology 274, H178–H186 (1998).

7. Peterson, E. C., Wang, Z. & Britz, G. Regulation of Cerebral Blood Flow. International Journal of Vascular Medicine 2011, 1–8 (2011).

8. Avery’s Diseases of the Newborn: [Edited by] Christine A. Gleason, Sherin U. Devaskar. (Elsevier/Saunders, Philadelphia, PA, 2012).

9. Jackson, W. F. Calcium-Dependent Ion Channels and the Regulation of Arteriolar Myogenic Tone. Front. Physiol. 12, 770450 (2021).

10. Davies, P. F. Flow-mediated endothelial mechanotransduction. Physiological Reviews 75, 519–560 (1995).

11. Klabunde, R. E. Cardiovascular Physiology Concepts. (Lippincott Williams & Wilkins/Wolters Kluwer, Philadelphia, PA, 2012).

12. Payne, S. J. Identifying the myogenic and metabolic components of cerebral autoregulation. Medical Engineering & Physics 58, 23–30 (2018).

13. Cipolla, M. J., McCall, A. L., Lessov, N. & Porter, J. M. Reperfusion Decreases Myogenic Reactivity and Alters Middle Cerebral Artery Function After Focal Cerebral Ischemia in Rats. Stroke 28, 176–180 (1997).

14. Cipolla, M. J. & Curry, A. B. Middle Cerebral Artery Function After Stroke: The Threshold Duration of Reperfusion for Myogenic Activity. Stroke 33, 2094–2099 (2002).

15. Niwa, K. et al. Cerebrovascular autoregulation is profoundly impaired in mice overexpressing amyloid precursor protein. American Journal of Physiology-Heart and Circulatory Physiology 283, H315–H323 (2002).

16. Panerai, R. B. Cerebral Autoregulation: From Models to Clinical Applications. Cardiovasc Eng 8, 42–59 (2008).

17. Klein, S. P., De Sloovere, V., Meyfroidt, G. & Depreitere, B. Dieerential Hemodynamic Response of Pial Arterioles Contributes to a Quadriphasic Cerebral Autoregulation Physiology. JAHA 11, e022943 (2022).

18. Ma, H. et al. Dynamic Cerebral Autoregulation in Embolic Stroke of Undetermined Source. Front. Physiol. 11, 557408 (2020).

19. Nogueira, R. C., Beishon, L., Bor-Seng-Shu, E., Panerai, R. B. & Robinson, T. G. Cerebral Autoregulation in Ischemic Stroke: From Pathophysiology to Clinical Concepts. Brain Sciences 11, 511 (2021).

20. Sherie, F. G. et al. A systematic review on the assessment of cerebral autoregulation in patients with Large Vessel Occlusion. Front Neurol 14, 1287873 (2023).

21. Vavilala, M. S. et al. Cerebral autoregulation in pediatric traumatic brain injury*: Pediatric Critical Care Medicine 5, 257–263 (2004).

22. Toth, P. et al. Traumatic brain injury-induced autoregulatory dysfunction and spreading depression-related neurovascular uncoupling: Pathomechanisms, perspectives, and therapeutic implications. Am J Physiol Heart Circ Physiol 311, H1118–H1131 (2016).

23. Novak, V., Novak, P., Spies, J. M. & Low, P. A. Autoregulation of Cerebral Blood Flow in Orthostatic Hypotension. Stroke 29, 104–111 (1998).

24. Claassen, J. A. & Zhang, R. Cerebral autoregulation in Alzheimer’s disease. J Cereb Blood Flow Metab 31, 1572–1577 (2011).

25. Wang, S. et al. Impact of impaired cerebral blood flow autoregulation on cognitive impairment. Front. Aging 3, 1077302 (2022).

26. Campbell, B. C. V. et al. Ischaemic stroke. Nat Rev Dis Primers 5, 70 (2019).

27. Kunz, A. & Iadecola, C. Chapter 14 Cerebral vascular dysregulation in the ischemic brain. in Handbook of Clinical Neurology vol. 92 283–305 (Elsevier, 2008).

28. Schwarz, S., Georgiadis, D., Aschoe, A. & Schwab, S. Eeects of Body Position on Intracranial Pressure and Cerebral Perfusion in Patients With Large Hemispheric Stroke. Stroke 33, 497–501 (2002).

29. Aries, M. J. H., Elting, J. W., De Keyser, J., Kremer, B. P. H. & Vroomen, P. C. A. J. Cerebral Autoregulation in Stroke: A Review of Transcranial Doppler Studies. Stroke 41, 2697–2704 (2010).

30. Jordan, J. D. & Powers, W. J. Cerebral Autoregulation and Acute Ischemic Stroke. Am J Hypertens 25, 946–950 (2012).

31. Castro, P., Azevedo, E. & Sorond, F. Cerebral Autoregulation in Stroke. Curr Atheroscler Rep 20, 37 (2018).

32. Gregori-Pla, C. et al. Blood flow response to orthostatic challenge identifies signatures of the failure of static cerebral autoregulation in patients with cerebrovascular disease. BMC Neurol 21, 154 (2021).

33. Reinhard, M. et al. Cerebral Autoregulation Dynamics in Acute Ischemic Stroke after rtPA Thrombolysis. Cerebrovasc Dis 26, 147–155 (2008).

34. Nogueira, R. C. et al. Cerebral autoregulation and response to intravenous thrombolysis for acute ischemic stroke. Sci Rep 10, 10554 (2020).

35. Meyer, M. et al. Impaired Cerebrovascular Autoregulation in Large Vessel Occlusive Stroke after Successful Mechanical Thrombectomy: A Prospective Cohort Study. Journal of Stroke and Cerebrovascular Diseases 29, 104596 (2020).

36. Lee, N. T. et al. Role of Purinergic Signalling in Endothelial Dysfunction and Thrombo-Inflammation in Ischaemic Stroke and Cerebral Small Vessel Disease. Biomolecules 11, 994 (2021).

37. Binder, N. F. et al. Leptomeningeal Collaterals Regulate Reperfusion in Ischemic Stroke. http://biorxiv.org/lookup/doi/10.1101/2023.02.25.529915 (2023) doi:10.1101/2023.02.25.529915.

38. García-Tornel, Á. et al. Leptomeningeal Collateral Flow Modifies Endovascular Treatment Efficacy on Large-Vessel Occlusion Strokes. Stroke 52, 299–303 (2021).

39. Fukuda, K. A. & Liebeskind, D. S. Evaluation of Collateral Circulation in Patients with Acute Ischemic Stroke. Radiologic Clinics of North America 61, 435–443 (2023).

40. Ursino, M. & Lodi, C. A. A simple mathematical model of the interaction between intracranial pressure and cerebral hemodynamics. Journal of Applied Physiology 82, 1256–1269 (1997).

41. Gao, E., Young, W. L., Pile-Spellman, J., Ornstein, E. & Ma, Q. Mathematical considerations for modeling cerebral blood flow autoregulation to systemic arterial pressure. American Journal of Physiology-Heart and Circulatory Physiology 274, H1023–H1031 (1998).

42. Ursino, M., Ter Minassian, A., Lodi, C. A. & Beydon, L. Cerebral hemodynamics during arterial and CO2 pressure changes: in vivo prediction by a mathematical model. American Journal of Physiology-Heart and Circulatory Physiology 279, H2439– H2455 (2000).

43. Aoi, M., Gremaud, P., Tran, H. T., Novak, V. & Olufsen, M. S. Modeling cerebral blood flow and regulation. in 2009 Annual International Conference of the IEEE Engineering in Medicine and Biology Society 5470–5473 (IEEE, Minneapolis, MN, 2009). doi:10.1109/IEMBS.2009.5334057.

44. Spronck, B., Martens, E. G. H. J., Gommer, E. D. & Van De Vosse, F. N. A lumped parameter model of cerebral blood flow control combining cerebral autoregulation and neurovascular coupling. American Journal of Physiology-Heart and Circulatory Physiology 303, H1143–H1153 (2012).

45. Lampe, R., Botkin, N., Turova, V., Blumenstein, T. & Alves-Pinto, A. Mathematical Modelling of Cerebral Blood Circulation and Cerebral Autoregulation: Towards Preventing Intracranial Hemorrhages in Preterm Newborns. Computational and Mathematical Methods in Medicine 2014, 1–9 (2014).

46. Henley, B. C., Shin, D. C., Zhang, R. & Marmarelis, V. Z. Compartmental and Data-Based Modeling of Cerebral Hemodynamics: Nonlinear Analysis. IEEE Trans Biomed Eng 64, 1078–1088 (2017).

47. Tong, Z., Catherall, M. & Payne, S. J. A multiscale model of cerebral autoregulation. Medical Engineering & Physics 95, 51–63 (2021).

48. Golubev, A., Kovtanyuk, A. & Lampe, R. Modeling of Cerebral Blood Flow Autoregulation Using Mathematical Control Theory. Mathematics 10, 2060 (2022).

49. Daher, A. & Payne, S. A network-based model of dynamic cerebral autoregulation. Microvascular Research 147, 104503 (2023).

50. Esfandi, H., Javidan, M., Anderson, R. M. & Pashaie, R. Depth-Dependent Contributions of Various Vascular Zones to Cerebral Autoregulation and Functional Hyperemia: An In-Silico Analysis. bioRxiv 2024.10.07.616950 (2024) doi:10.1101/2024.10.07.616950.

51. Epp, R. et al. The role of leptomeningeal collaterals in redistributing blood flow during stroke. PLoS Comput Biol 19, e1011496 (2023).

52. Adams, M. D., Winder, A. T., Blinder, P. & Drew, P. J. The pial vasculature of the mouse develops according to a sensory-independent program. Sci Rep 8, 9860 (2018).

53. Epp, R., Schmid, F. & Jenny, P. Fast convergence strategy for ambiguous inverse problems based on hierarchical regularization. Journal of Computational Physics 489, 112264 (2023).

54. Boas, D. A., Jones, S. R., Devor, A., Huppert, T. J. & Dale, A. M. A vascular anatomical network model of the spatio-temporal response to brain activation. NeuroImage 40, 1116–1129 (2008).

55. Schmid, F., Reichold, J., Weber, B. & Jenny, P. The impact of capillary dilation on the distribution of red blood cells in artificial networks. American Journal of Physiology-Heart and Circulatory Physiology 308, H733–H742 (2015).

56. Secomb, T. W. Blood Flow in the Microcirculation. Annu. Rev. Fluid Mech. 49, 443–461 (2017).

57. Schmid, F., Tsai, P. S., Kleinfeld, D., Jenny, P. & Weber, B. Depth-dependent flow and pressure characteristics in cortical microvascular networks. PLoS Comput Biol 13, e1005392 (2017).

58. Schmid, F., Barrett, M. J. P., Obrist, D., Weber, B. & Jenny, P. Red blood cells stabilize flow in brain microvascular networks. PLoS Comput Biol 15, e1007231 (2019).

59. Epp, R., Schmid, F., Weber, B. & Jenny, P. Predicting Vessel Diameter Changes to Up-Regulate Biphasic Blood Flow During Activation in Realistic Microvascular Networks. Front. Physiol. 11, 566303 (2020).

60. Schmid, F., Conti, G., Jenny, P. & Weber, B. The severity of microstrokes depends on local vascular topology and baseline perfusion. eLife 10, e60208 (2021).

61. Pries, A. R. & Secomb, T. W. Microvascular blood viscosity in vivo and the endothelial surface layer. American Journal of Physiology-Heart and Circulatory Physiology 289, H2657–H2664 (2005).

62. Absi, R. Revisiting the pressure-area relation for the flow in elastic tubes: Application to arterial vessels. Series on Biomechanics 32, pp.47–59 (2018).

63. Rammos, K. A computer model for the prediction of left epicardial coronary blood flow in normal, stenotic and bypassed coronary arteries, by single or sequential grafting. Cardiovascular Surgery 6, 635–648 (1998).

64. Sherwin, S. J., Franke, V., Peiró, J. & Parker, K. One-dimensional modelling of a vascular network in space-time variables. Journal of Engineering Mathematics 47, 217–250 (2003).

65. Urquiza, S. A., Blanco, P. J., Vénere, M. J. & Feijóo, R. A. Multidimensional modelling for the carotid artery blood flow. Computer Methods in Applied Mechanics and Engineering 195, 4002–4017 (2006).

66. Ranjan, V., Xiao, Z. & Diamond, S. L. Constitutive NOS expression in cultured endothelial cells is elevated by fluid shear stress. American Journal of Physiology-Heart and Circulatory Physiology 269, H550–H555 (1995).

67. Chiu, J. J., Wung, B. S., Hsieh, H. J., Lo, L. W. & Wang, D. L. Nitric Oxide Regulates Shear Stress–Induced Early Growth Response-1: Expression via the Extracellular Signal–Regulated Kinase Pathway in Endothelial Cells. Circulation Research 85, 238–246 (1999).

68. Ashpole, N. E., Overby, D. R., Ethier, C. R. & Stamer, W. D. Shear stress-triggered nitric oxide release from Schlemm’s canal cells. Invest Ophthalmol Vis Sci 55, 8067–8076 (2014).

69. Koide, M., Ferris, H. R., Nelson, M. T. & Wellman, G. C. Impaired Cerebral Autoregulation After Subarachnoid Hemorrhage: A Quantitative Assessment Using a Mouse Model. Front. Physiol. 12, 688468 (2021).

70. Glück, C. et al. Pia-FLOW: Deciphering hemodynamic maps of the pial vascular connectome and its response to arterial occlusion. Proc. Natl. Acad. Sci. U.S.A. 121, e2402624121 (2024).

71. Glandorf, L. et al. In Vivo Network-Level Cerebrovascular Mapping Reveals the Impact of Flow Topology on Capillary Stalls After Stroke. Preprint at 10.1101/2025.07.28.667165 (2025).

72. Loredo, J. Sleep quality and blood pressure dipping in obstructive sleep apnea. American Journal of Hypertension 14, 887–892 (2001).

73. Friedman, O. & Logan, A. G. Nocturnal blood pressure profiles among normotensive, controlled hypertensive and refractory hypertensive subjects. Canadian Journal of Cardiology 25, S312–S316 (2009).

74. Koike, M. A., Green, K. N., Blurton-Jones, M. & LaFerla, F. M. Oligemic Hypoperfusion Diffrentially Affcts Tau and Amyloid-β. The American Journal of Pathology 177, 300–310 (2010).

75. ElAli, A., Thériault, P., Préfontaine, P. & Rivest, S. Mild chronic cerebral hypoperfusion induces neurovascular dysfunction, triggering peripheral beta-amyloid brain entry and aggregation. acta neuropathol commun 1, 75 (2013).

76. Xie, L. et al. Sleep Drives Metabolite Clearance from the Adult Brain. Science 342, 373–377 (2013).

77. Iliyasu, M. O., Musa, S. A., Oladele, S. B. & Iliya, A. I. Amyloid-beta aggregation implicates multiple pathways in Alzheimer’s disease: Understanding the mechanisms. Front. Neurosci. 17, 1081938 (2023).

78. Pradeepkiran, J. A., Baig, J., Islam, M. A., Kshirsagar, S. & Reddy, P. H. Amyloid-β and Phosphorylated Tau are the Key Biomarkers and Predictors of Alzheimer’s Disease. Aging and disease 16, 658 (2025).

79. Pluta, R., Czuczwar, S. J., Januszewski, S. & Jabłoński, M. The Many Faces of Post-Ischemic Tau Protein in Brain Neurodegeneration of the Alzheimer’s Disease Type. Cells 10, 2213 (2021).

80. Rost, N. S. et al. Post-Stroke Cognitive Impairment and Dementia. Circulation Research 130, 1252–1271 (2022).

81. Khan, R., Devlin, P., Urayama, A. & Ritzel, R. M. Models and mechanisms of post-stroke dementia and cognitive impairment. Front. Stroke 4, 1563924 (2025).

82. Toth, P., Tarantini, S., Csiszar, A. & Ungvari, Z. Functional vascular contributions to cognitive impairment and dementia: mechanisms and consequences of cerebral autoregulatory dysfunction, endothelial impairment, and neurovascular uncoupling in aging. American Journal of Physiology-Heart and Circulatory Physiology 312, H1– H20 (2017).

83. Miller, L. R. et al. IGF1R deficiency in vascular smooth muscle cells impairs myogenic autoregulation and cognition in mice. Front. Aging Neurosci. 16, 1320808 (2024).

84. Fjorbak, C. L., Kutuzov, N. P., Groves, T., Lauritzen, M. & Grubb, S. Brain precapillary sphincters modulate myogenic tone in adult and aged mice. GeroScience (2025) doi:10.1007/s11357-025-01720-8.

85. Salinet, A. S. M., Robinson, T. G. & Panerai, R. B. Cerebral blood flow response to neural activation after acute ischemic stroke: a failure of myogenic regulation? J Neurol 260, 2588–2595 (2013).

86. Kim, B. J., Singh, N. & Menon, B. K. Hemodynamics of Leptomeningeal Collaterals after Large Vessel Occlusion and Blood Pressure Management with Endovascular Treatment. J Stroke 23, 343–357 (2021).

87. Litman, M., Martin, K., Spratt, N. J. & Beard, D. J. Quantification of leptomeningeal collateral blood flow in hypertensive rats during ischemic stroke. Journal of Stroke and Cerebrovascular Diseases 34, 108195 (2025).

88. Faber, J. E. Collateral blood vessels in stroke and ischemic disease: Formation, physiology, rarefaction, remodeling. J Cereb Blood Flow Metab 45, 1007–1030 (2025).

89. Palomares, S. & Cipolla, M. Myogenic Tone as a Therapeutic Target for Ischemic Stroke. CVP 12, 788–800 (2014).

90. Coucha, M., Li, W., Johnson, M. H., Fagan, S. C. & Ergul, A. Protein nitration impairs the myogenic tone of rat middle cerebral arteries in both ischemic and nonischemic hemispheres after ischemic stroke. American Journal of Physiology-Heart and Circulatory Physiology 305, H1726–H1735 (2013).

91. Nag, S. & Kilty, D. W. Cerebrovascular Changes in Chronic Hypertension: Protective Effects of Enalapril in Rats. Stroke 28, 1028–1034 (1997).

92. Alves, L., Hashiguchi, D., Loss, C. M., Van Praag, H. & Longo, B. M. Vascular dysfunction in Alzheimer’s disease: Exploring the potential of aerobic and resistance exercises as therapeutic strategies. Journal of Alzheimer’s Disease 104, 963–979 (2025).

93. Ren, Y. et al. Enhanced myogenic response in the afferent arteriole of spontaneously hypertensive rats. Am J Physiol Heart Circ Physiol 298, H1769–1775 (2010).

94. Tzeng, Y.-C. & Ainslie, P. N. Blood pressure regulation IX: cerebral autoregulation under blood pressure challenges. Eur J Appl Physiol 114, 545–559 (2014).

95. Chalothorn, D., Clayton, J. A., Zhang, H., Pomp, D. & Faber, J. E. Collateral density, remodeling, and VEGF-A expression differ widely between mouse strains. Physiological Genomics 30, 179–191 (2007).

96. Taherzadeh, Z. et al. Strain-dependent susceptibility for hypertension in mice resides in the natural killer gene complex. American Journal of Physiology-Heart and Circulatory Physiology 298, H1273–H1282 (2010).

## Reference

1. Campolongo, F., Cariboni, J. & Saltelli, A. An effective screening design for sensitivity analysis of large models. Environmental Modelling & Software 22, 1509–1518 (2007).

2. Global Sensitivity Analysis: The Primer. (John Wiley, Chichester, England Hoboken, NJ, 2008). doi:10.1002/9780470725184.

3. Daher, A. & Payne, S. A dynamic multiscale model of cerebral blood flow and autoregulation in the microvasculature. Applied Mathematical Modelling 123, 213–240 (2023).

