## Supplementary Material for "Loss of Cerebral Autoregulation After Stroke Drives Abnormal Perfusion Patterns"

### *Supplementary Figures*

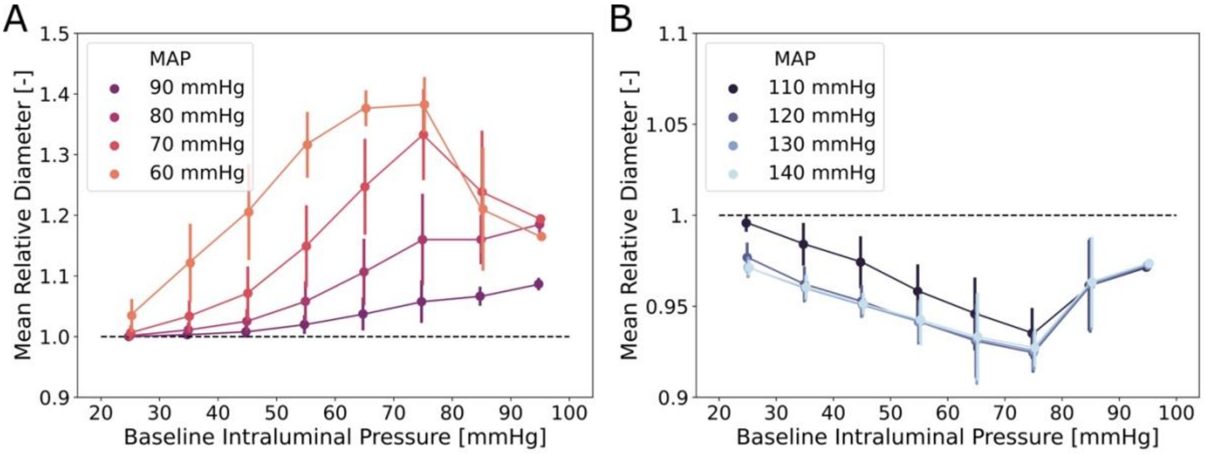


**Figure S1. (A)** Mean relative vessel diameter as a function of baseline intraluminal pressure under different levels of mean arterial pressure (MAP) reduction (60–90 mmHg). **(B)** Mean relative vessel diameter as a function of baseline intraluminal pressure under increasing MAP levels (110–140 mmHg). The baseline MAP is 100 mmHg. Results represent the mean ± standard deviation (error bars) computed across all arteries of all four microvascular networks.

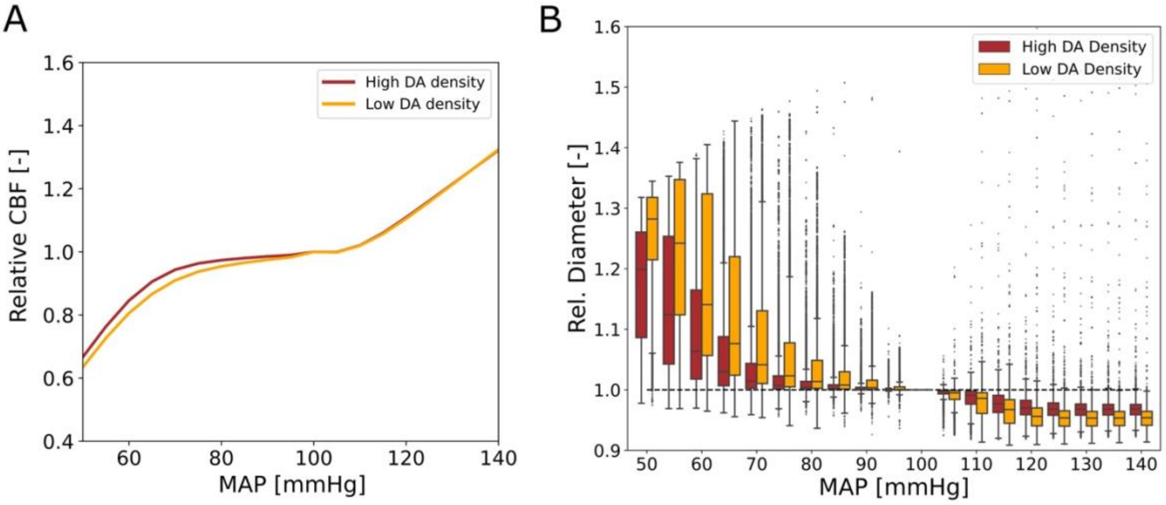


**Figure S2. (A)** Mean relative cerebral blood flow (CBF) as a function of mean arterial pressure (MAP) for microvascular networks with high and low descending arteriole (DA) densities. **(B)** Distribution of relative arterial diameters across all vessels in the two configurations, shown as a function of MAP. All values are reported relative to their respective baseline values at 100 mmHg MAP.

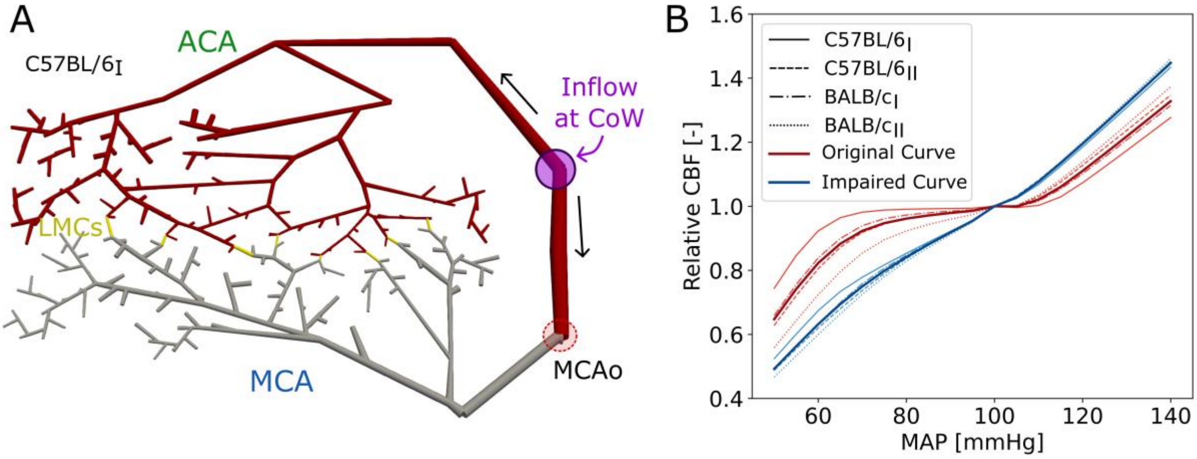


**Figure S3. (A)** Microvascular network (C57BL/6$_{\mathrm{I}}$) illustrating the distribution of passive arteries (gray) and actively regulating vessels (red). The site of the previously simulated middle cerebral artery occlusion (MCAo) is indicated by the dashed red circle. Autoregulatory impairment extends from the MCAo location throughout the MCA territory, reaching up to the leptomeningeal collaterals (LMCs, shown in yellow). This configuration represents the fully impaired case. **(B)** Relative cerebral blood flow (CBF) normalized to baseline values as a function of mean arterial pressure (MAP), comparing fully regulated (original) and fully impaired conditions for each of the four networks. The two darker-colored lines represent the average curves across all networks for each condition.

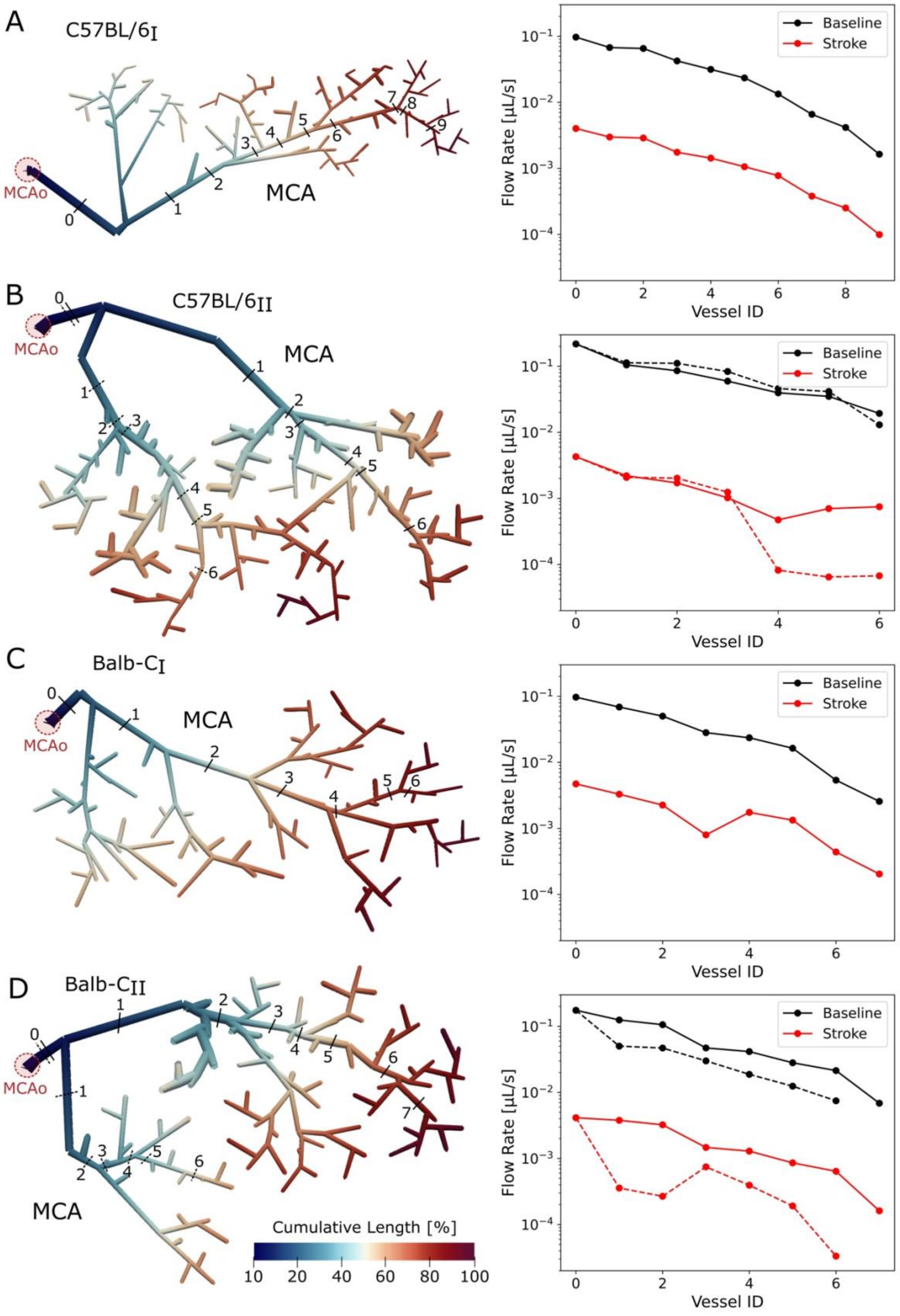


**Figure S4.** Visualization of cumulative vessel length and corresponding flow rates under baseline and stroke conditions. (A–D) Left: Middle cerebral artery (MCA)-sided vasculature for each microvascular network (C57BL/6$_{I}$, C57BL/6$_{II}$, Balb-C$_{I}$, and Balb-C$_{II}$), colored by cumulative vessel length, normalized to the maximum value within each network. Selected vessel segments are labeled based on their cumulative distance from the middle cerebral artery occlusion (MCAo, red circle). Right: Corresponding flow rates [μL/s] in each labeled segment under baseline (black) and stroke (red) conditions, illustrating the progressive reduction in perfusion along the vascular path after MCAo.

### *Supplementary Tables*

**Table S1.** Parameters used in the autoregulation model and corresponding structural characteristics of the microvasculature. References for parameter values are indicated.

| Parameter | Parameter Value | Literature |
|---|---|---|
| ***Autoregulation model parameters*** | | |
| Myogenic Sensitivity ($S\sigma$) | 4.0 | [1] |
| Endothelial Sensitivity ($S\tau$) | 0.5 | [1] |
| Sigmoidal Slope ($G$) | 0.1 | [1] |
| Maximum positive compliance changes ($\Delta C^{+}/C_0$) | 10 | [1] |
| Maximum negative compliance changes ($\Delta C^{-}/C_0$) | 0.8 | [1] |
| ***Characteristics of microvasculature*** | | |
| Young's Modulus ($E$) [Pa] | | |
| Arteries (radius ≥ 20 µm) | $6.30 \times 10^5$ | [2] |
| Arteries (15 µm ≤ radius < 20 µm) | $2.60 \times 10^5$ | [2] |
| Arteries (radius < 15 µm) | $1.60 \times 10^5$ | [2] |
| Veins | $3.88 \times 10^5$ | [2] |
| Capillaries | $3.70 \times 10^5$ | [2] |
| Vessel wall thickness to diameter ratio ($h/D$) | | |
| Arteries | 0.2 | [2] |
| Veins & Capillaries | 0.1 | [3] |

**Table S2.** Relative diameter changes of leptomeningeal collaterals (LMCs) at stroke conditions, normalized to baseline values (prior to stroke) for each microvascular network (MVN). Mean ± standard deviation (SD) across all networks is also shown.

| MVNs | Relative diameter |
|---|---|
| C57BL/6$_{\mathrm{I}}$ | |
| | 1.15 |
| | 1.15 |
| | 1.16 |
| | 1.16 |
| | 1.15 |
| | 1.19 |
| | 1.17 |
| | 1.16 |
| C57BL/6$_{\mathrm{II}}$ | |
| | 1.25 |
| | 1.23 |
| | 1.25 |
| | 1.22 |
| Balb-C$_{\mathrm{I}}$ | |
| | 1.28 |
| | 1.27 |
| | 1.27 |
| | 1.27 |
| | 1.27 |
| | 1.26 |
| | 1.24 |
| Balb-C$_{\mathrm{II}}$ | |
| | 1.34 |
| | 1.26 |
| | 1.32 |
| | 1.26 |
| Mean ± SD | 1.23 ± 0.06 |
| Literature[4] | 1.12 ± 0.14 |

### Reference


1. Daher, A. & Payne, S. A network-based model of dynamic cerebral autoregulation. *Microvascular Research* **147**, 104503 (2023).
2. Salotto, A. G., Muscarella, L. F., Melbin, J., Li, J. K.-J. & Noordergraaf, A. Pressure pulse transmission into vascular beds. *Microvascular Research* **32**, 152–163 (1986).
3. Enouri, S., Monteith, G. & Johnson, R. Characteristics of myogenic reactivity in isolated rat mesenteric veins. *American Journal of Physiology-Regulatory, Integrative and Comparative Physiology* **300**, R470–R478 (2011).
4. Glück, C. *et al.* Pia-FLOW: Deciphering hemodynamic maps of the pial vascular connectome and its response to arterial occlusion. *Proc. Natl. Acad. Sci. U.S.A.* **121**, e2402624121 (2024).
